# Theory of transcription bursting: Stochasticity in the transcription rates

**DOI:** 10.1101/2019.12.18.880435

**Authors:** Rajamanickam Murugan

## Abstract

Transcription bursting creates variation among the individuals of a given population. Bursting emerges as the consequence of turning on and off the transcription process randomly. There are at least three sub-processes involved in the bursting phenomenon with different timescale regimes viz. flipping across the on-off state channels, microscopic transcription elongation events and the mesoscopic transcription dynamics along with the mRNA recycling. We demonstrate that when the flipping dynamics is coupled with the microscopic elongation events, then the distribution of the resultant transcription rates will be over-dispersed. This in turn reflects as the transcription bursting with over-dispersed non-Poisson type distribution of mRNA numbers. We further show that there exist optimum flipping rates (*α_C_*, *β_C_*) at which the stationary state Fano factor and variance associated with the mRNA numbers attain maxima. These optimum points are connected via 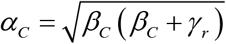. Here *α* is the rate of flipping from the on-state to the off-state, *β* is the rate of flipping from the off-state to the on-state and *γ_r_* is the decay rate of mRNA. When *α* = *β* = *χ* with zero rate in the off-state channel, then there exist optimum flipping rates at which the non-stationary Fano factor and variance attain maxima. Here 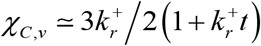 (here 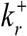 is the rate of transcription purely through the on-state elongation channel) is the optimum flipping rate at which the variance of mRNA attains a maximum and *χ*_*C*, *κ*_ ≃ 1.72/*t* is the optimum flipping rate at which the Fano factor attains a maximum. Close observation of the transcription mechanism reveals that the RNA polymerase performs several rounds of stall-continue type dynamics before generating a complete mRNA. Based on this observation, we model the transcription event as a stochastic trajectory of the transcription machinery across these on-off state elongation channels. Each mRNA transcript follows different trajectory. The total time taken by a given trajectory is the first passage time (FPT). Inverse of this FPT is the resultant transcription rate associated with the particular mRNA. Therefore, the time required to generate a given mRNA transcript will be a random variable. For a stall-continue type dynamics of RNA polymerase, we show that the overall average transcription rate can be expressed as 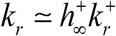 where 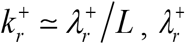 is the microscopic transcription elongation rate in the on-state channel and *L* is the length of a complete mRNA transcript and 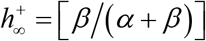 is the stationary state probability of finding the transcription machinery in the on-state.

## INTRODUCTION

Subcellular process such as transcription and translation of various genes are essential for the survival of an organism [1, 2]. Since these processes are mesoscopic in nature, their outcomes such as mRNAs and protein molecules are prone to great number fluctuations. Particularly, cells with identical genetic materials, produce different levels of mRNAs and proteins corresponding to various genes at a given time point. Such variations in the expression of different genes across the population of individual cells is essential for the survival of an organism against various extreme environmental conditions. The molecular number fluctuations across various cells of a population, can be influenced by both intrinsic and extrinsic factors. The intrinsic noise in gene expression arises well within the system itself which is characterized by a set of kinetic parameters such as transcription and translation rate constants. On the other hand, external environmental factors such as temperature and nutrient fluctuations can perturb these intrinsic kinetic parameters which ultimately emerges as the extrinsic noise component [3].

The stochasticity of the constitutive gene expression has been extensively investigated both theoretically and experimentally [4–10]. The fluctuations in the number of mRNAs and protein molecules can be well characterized by the corresponding population *mean*, *variance*, *coefficient of variation* and the *Fano factor*. Here *coefficient of variation* = (*standard deviation* / *mean*) and *Fano factor* = (*variance* / *mean*). The Fano factor measures the extent of deviation of the molecular number fluctuations from the standard Poisson process for which the Fano factor = 1. Populations which exhibit Fano factor < 1 are under-dispersed and those populations which exhibit Fano factor > 1 are over-dispersed. Detailed theoretical calculations and subsequent experimental studies on the unregulated gene expression have shown that the Fano factors associated with the protein number fluctuations is more than one and its deviation from the Poisson increases linearly with the translational efficiency [8, 11]. Here the *translational efficiency* = *translation rate* / *decay rate of mRNA molecules*. On the other hand, the molecular number fluctuations in mRNAs follows a typical Poisson process with Fano factor = 1.

Gene expression was initially thought as a continuous process and the probability density functions associated with the mRNAs and protein number fluctuations are assumed to be a monomodal type. Later experiments revealed the interrupted and bursting nature of mRNAs and protein numbers along the temporal axis [12, 13]. Such transcriptional bursting can result in a bimodal or multimodal type density functions associated with the number of mRNAs and proteins [12, 14, 15]. By definition, two stage gene expression involves only transcription and translation processes and three stage gene expression incudes on-off state dynamics of the promoter along with the transcription and translation. The main sources of transcription bursting can be attributed to the initiation and elongation steps [14]. In the process of transcription initiation, the promoter seems to be turned on and off in a random manner via the binding-unbinding of the regulatory transcription factors (TFs). On the other hand, the RNA polymerase enzyme complex (RNAP) can undergo stall-continue type dynamics in the transcription elongation process. Both these types of dynamics ultimately introduce a time dependent stochasticity in the overall transcription rate constant. Transcription bursting phenomenon has been studied in detail [16–23]. The effects of various factors such as negative feedback [17], presence of enhancer elements [24, 25] on the frequency of bursting and burst size have also been unraveled in detail.

Both the binding-unbinding of regulatory TFs proteins with the promoter and stall-continue type dynamics of RNAP in the elongation process ultimately switch on or off the transcription event in a time dependent and random manner. Generally, transcription elongation generates positive supercoiling ahead of the RNAP complex and leaves negative supercoil trail behind the RNAP [26–28]. For a smooth transcription elongation process, these positive and negative supercoils need to be relaxed by the gyrase and topoisomerase I enzymes respectively [14]. When there is a lack in the number of gyrase molecules to handle the accumulation of positive supercoils ahead of the transcribing RNAP, then RNAP complex cannot progress further and subsequently transcription event will be stalled. When enough gyrase molecules arrive and relax the accumulated positive supercoils, then the transcription event continues and so on. Recent studies revealed the possibility of co-transcription of several RNAPs in a convoy with finite interval among them [29, 30]. Within a convoy of RNAPs with appropriate spacing among them, there is a possibility that the negative supercoil trail of the first RNAP will be cancelled by the positive supercoil ahead of the second RNAP and so on. This is the fluid model of transcription elongation which requires generally short distances among the RNAPs [29]. The cooperative interactions among the RNAPs of a convoy can be either collaborative or antagonistic depending on the spacing among them which can be well modelled by the push and push-pull mechanisms [29–31]. The overall interactions among the RNAPs of a convoy ultimately result in the random speed up or slowdown of the transcription elongation dynamics of individual RNAPs apart from the stall-continue type dynamics of the convoy in a time dependent manner.

Clearly, there are three different timescale regimes involved in the transcription bursting viz. (**a**) the timescale associated with the microscopic elongation transitions in the generation of a complete mRNA transcript, (**b**) the timescale associated with the on-off flipping dynamics and (**c**) the timescale associated with the mesoscopic dynamics of mRNA along with its recycling. Interplay of these processes (**a**), (**b**), and (**c**) results in the over-dispersion of the distribution of mRNA numbers. Almost all the theoretical and experimental studies on the transcription bursting 1) assumed slower timescale for the on-off state flipping than the timescale of transcription and decay and 2) concentrated on the steady state of the gene expression. In this paper, using a combination of theoretical and simulation tools we will show that the flipping across the on-off transcription elongation channels introduce stochasticity in the transcription rate and there exists an optimum flipping rate at which the variance and Fano factor of mRNA numbers attain the maxima.

## THEORY

Let us consider the typical transcription event along with the recycling of mRNAs well within the cellular environment as depicted in **Fig. 1**. When the transcription happens in a “stall-continue” or “on-off” mode of RNA polymerase enzyme complex (RNAP in prokaryotes and RNA Pol II in eukaryotes), then it can be described by the following set of coupled master equations.

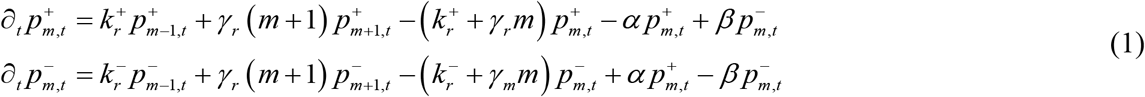

**FIGURE 1.**
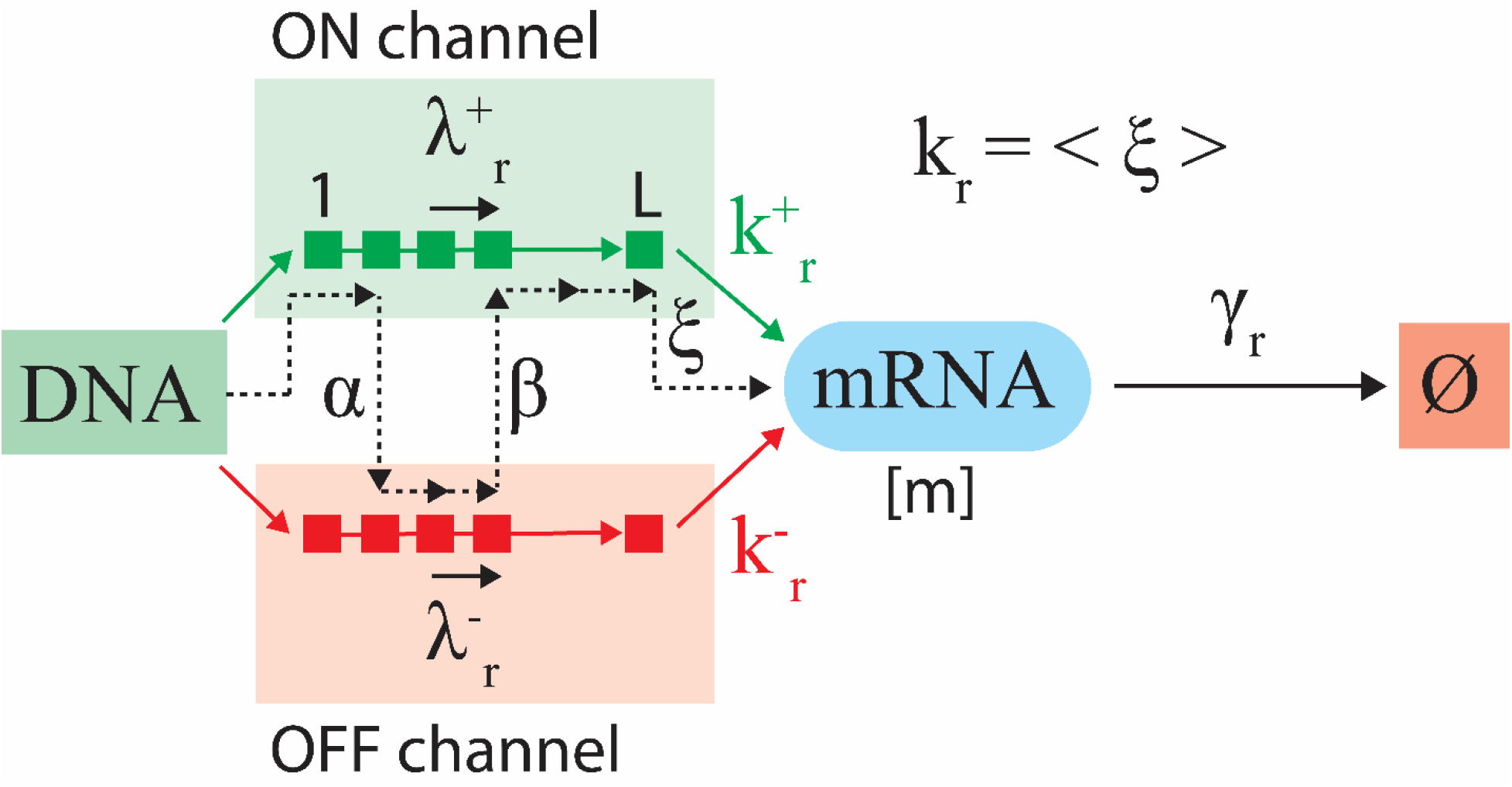
Stochastic transcription rate and transcription bursting. In this generalized model, the mRNA transcript of size *L* bp can be generated via either pure on or off state channels with rates 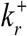 and 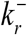 respectively. The microscopic transition rates 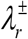 characterize the addition of individual nucleotides at the end of growing mRNA in the process of transcription elongation. The resultant transcription rate will be the inverse of the first passage time required to generate a complete mRNA. When there is a flipping across these on-off states then the resultant transcription rate *ξ* fluctuate across 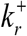 and 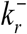 in a random manner where 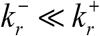 in general. The ensemble average of *ξ* across several trajectories of mRNA synthesis will be the average transcription rate *k*_*r*_ = ⟨*ξ*⟩. Here (+) denote the on-state and (−) denotes the off-state of transcription and *γ*_*r*_ is the decay rate constant of mRNAs. The rate of flipping from the on-state to the off state is *α* and the rate of flipping from the off-state to the on-state in *β*. The mesoscopic transcription rates 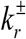 are connected to the microscopic ones 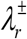 via the relationship 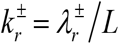. As a result, flipping across on-off states, the overall effective transcription rate *k_r_* will be somewhere in between 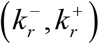. Here the dotted line is a stochastic transcription trajectory which varies from transcript to transcript. This means that the effective transcription rate *ξ* (it is not a constant anymore) varies from transcript to transcript and it is a stochastic quantity.

Let us denote *p_m,t_* as the overall probability of finding *m* number of mRNA molecules at time *t* starting from zero number of mRNAs at time *t* = 0. The probabilities of finding the transcription machinery at the on-off states are described by 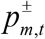 where the superscript (+) denotes the on-state and (−) denotes the off-state and the corresponding transcription rates are 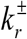 (s^−1^ when mRNA concentration is measured in numbers or M/s when mRNA concentration is measured in M). Let us assume that the length of the complete mRNA transcript is *L* bp where 1 bp = 3.4 × 10^−10^ m. When there is a flipping across the on-off state channels, then the synthesis of a given mRNA transcript of interest will eventually follow a random trajectory as described in **Fig. 1**. We denote the resultant transcription rate of this random trajectory as *ξ* which is the inverse of the total time taken by the particular trajectory and it varies from trajectory to trajectory. We define the *average effective or resultant transcription rate* as *k_r_* which is the average of *ξ* across several trajectories of the transcription event as *k*_*r*_ = ⟨*ξ*⟩. In general, one finds that 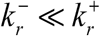 and 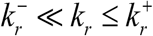.

We have assumed a non-zero transcription rate 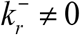 for the off-state channel mainly due to the fact that the RNA polymerase complex completely stalls only when the level of positive supercoiling accumulating ahead of transcription is strong enough to completely stop its further progression. Otherwise, the progression of the RNA polymerase over transcription will be slower than the normal speed (but not equal to zero). Here *α* (s^−1^) is the rate of transition from the on-state to the off-state and *β* (s^−1^) is the rate of transition from the off-state to the on-state of transcription. The ratio *σ* = *k_r_* / *α* is generally defined as the *transcription efficiency* [3] or the burst size. Here *σ* is the average number of mRNA molecules generated in the on-state of transcription. The first order decay rate of mRNAs is described by *γ*_*r*_ (s^−1^). The main assumption in **Eqs. 1** is that the timescales associated with the flipping rates (*α*, *β*) are comparable with or higher than the timescales associated with the transcription rates 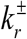. We will show in the later sections that 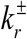 will be rescaled with (*α*, *β*) when the timescales associated with (*α*, *β*) are shorter than the timescales associated with the generation a full mRNA transcript. When the average transcription speed of the RNAP is 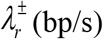 and the length of the final mRNA transcript is *L* bp, then the average transcription rate through the respective on and off channels will be 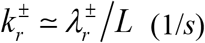. In other words, 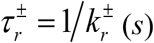 is the average time required to generate a full mRNA transcript via the respective on-off channels. Here one should note that the maximum elongation of speed of RNA pol II seems to be close to ~100 bp/s [32, 33]. Typical transcription elongation speed of the prokaryotic RNAP complex is 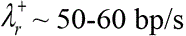 [2]. The average resultant transcription rate *k_r_* will be a function of the variables 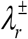, *L*, *α* and *β*. The initial and normalization conditions associated with **Eqs. 1** can be given as follows.

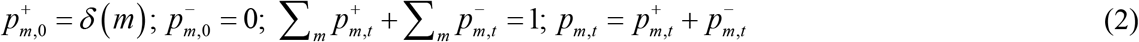

**Eqs. 1** and **2** can be solved using the standard method of generating function formalism. Upon defining the generating functions as 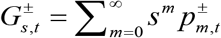, **Eq. 1** can written as follows.

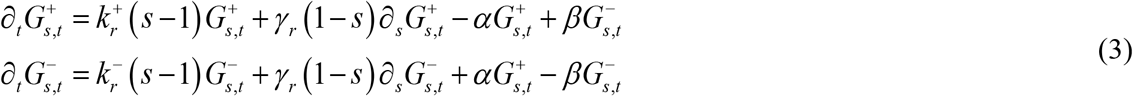

General solution to **Eqs. 3** was obtained in terms of confluent hypergeometric functions in Ref. [34] using Galilean type transformation of variables (Eqs. 13 and 14 in Ref. 31). However, these solutions in terms of confluent hypergeometric functions are very difficult to handle. Particularly, the derivation of the expressions for the variance and the Fano factor associated with the mRNA number fluctuations will not be a difficult task. To avoid this hurdle, we follow the route of case-splitting technique to derive all the possible approximations under various conditions which are simple and also easy to handle. Upon performing few basic operations, this system can be rewritten as the following set of two uncoupled partial differential equations.

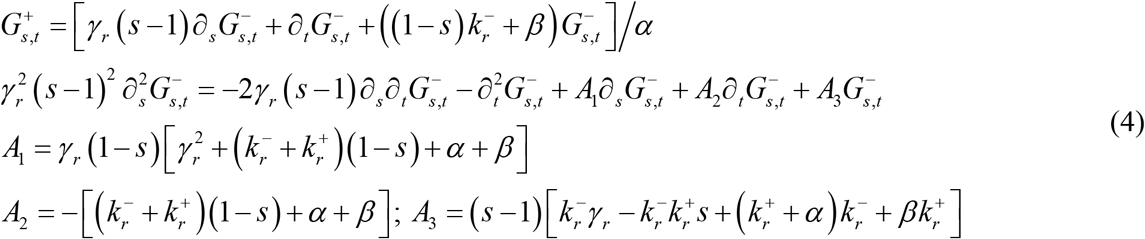

Here we have the initial conditions as 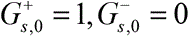 [35]. **Eqs. 3**-**4** are the central equations of this paper from which we derive several interesting results. Upon approximately solving these partial differential equations, one can recover the expressions for the probability density functions associated with the mRNA fluctuations 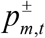 as follows.

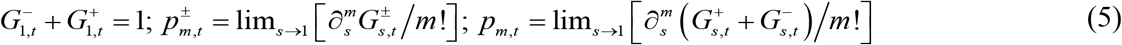

Further, using these generating functions, one can straightforwardly derive the expressions for the various time dependent statistical quantities associated with the population of mRNAs such as mean (*η*_*m,t*_), variance (*v_m,t_*), coefficient of variation (*μ_m,t_*) and Fano factor (*κ*_*m,t*_) as follows.

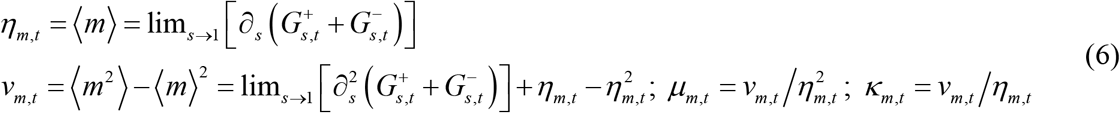

With this background, we will consider the following exactly solvable cases and other interesting approximations of **Eqs. 3**-**4**.

**Case I**. *α* ≠ *β*; 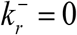; *γ*_*r*_ = 0

This case has been investigated earlier in Refs. [35, 36] (Eqs. 11 and 12 in Ref. [35]). Under these conditions, **Eqs. 4** can be reduced to the following form which is exactly solvable.

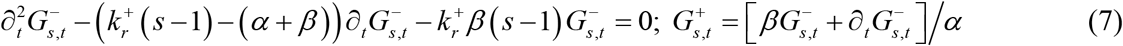

The general solution to this set of partial differential equations can be written as follows.

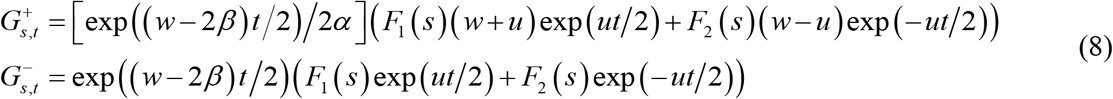

Here 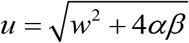 where 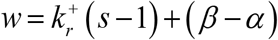. Noting that 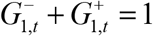, the functions *F*_1_(*s*) and *F*_2_(*s*) and the particular solution to **Eqs. 8** can be derived as follows.

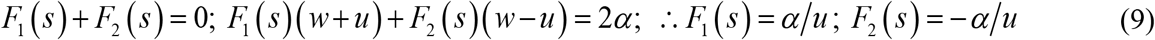

Upon substituting these functions into **Eqs. 8** one finally obtains the following expression for the generating functions.

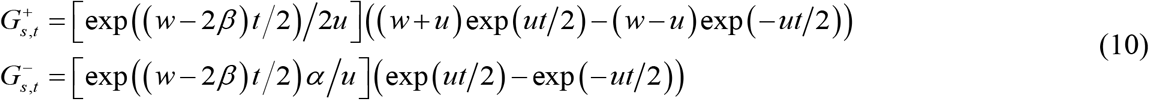

Using the overall generating function, one can derive the mean number of mRNAs as follows.

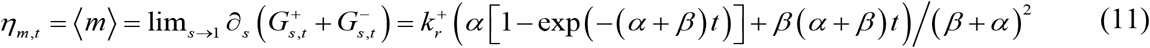

For short timescales, 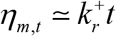 which is similar to the deterministic result and for long timescale it behaves as 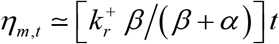. Noting that 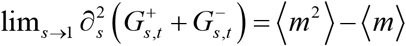, which can be written explicitly as follows.

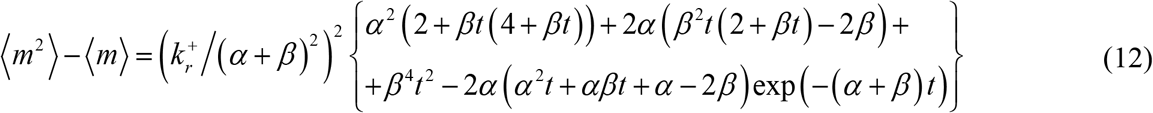

Using **Eqs. 11** and **12** one can write down the variance, coefficient of variation and Fano factor of mRNAs as follows.

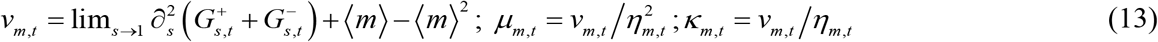

It is remarkable to note down the following limits as time tends towards infinity.

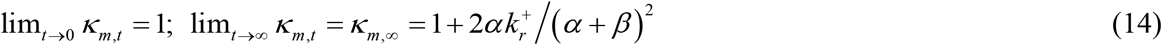

Clearly, in **Eqs. 14** the stationary state Fano factor *κ*_*m*,∞_ attains a maximum value at an optimum *α*_*C*_ = *β* which can be obtained by solving [*∂κ*_*m*,∞_ / *∂α*] = 0 for *α*.

**Case II.** *α* = *β* = *χ*; 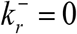; *γ*_*r*_ ≠ 0

In this case, **Eqs. 4** reduce to the following set of uncoupled partial differential equations.

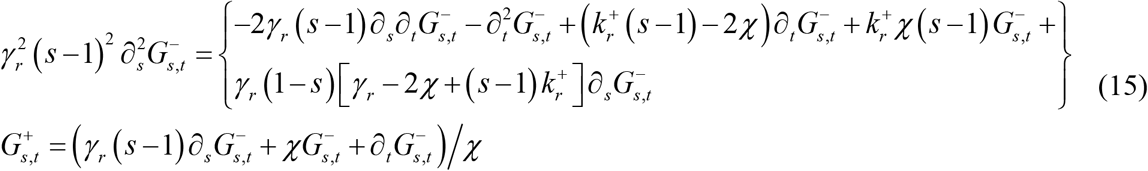

Similar to this case with settings (*α* ≠ *β*; 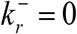; *γ*_*r*_ ≠ 0) was considered in Refs. [35, 36]. The first term of the series expansion was obtained using recurrence relation method in Ref. [35]. Interestingly, the solution set for these conditions was obtained in terms of confluent hypergeometric functions using Galilean type transformation of variables in Ref. [36] (Eqs. A2-A5 in Ref. [36]). Remarkably, when *χ* → ∞, then **Eqs. 15** reduce to the following form.

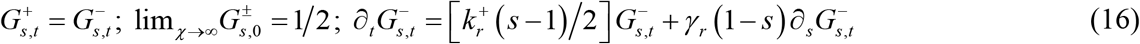

These equations are exactly solvable as follows.

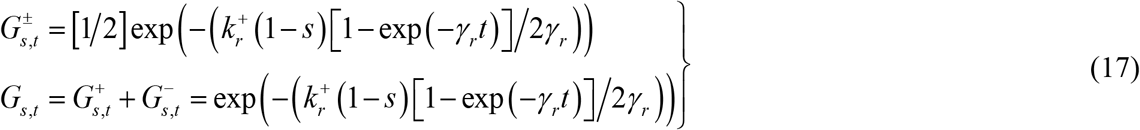

Using these generating functions, one can derive the Poisson type expressions for the probability density function associated with the mRNA population as follows.

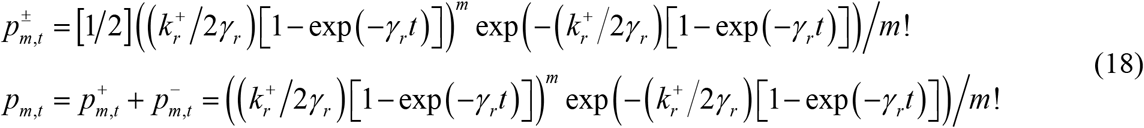

Various statistical properties of mRNA population can be derived as follows.

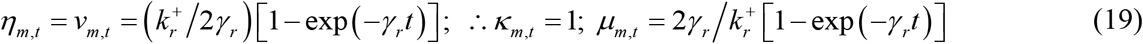

These results clearly suggest that as *χ* → ∞, the overall average or effective transcription rate *k_r_* rescales from 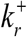 to 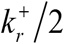. The condition χ → ∞ implies that the system switches infinite number of times between on and off states with equal amount of dwell times at both these states. That is to say, the timescale of flipping between on-off states are comparable with the timescales of the microscopic transcription elongation events. Therefore, the condition *χ* → ∞ violates the assumption of **Eqs. 1** and **Eqs. 16**-**18** since the transcription rates will be a function of the flipping rate *χ* under such conditions.

**Case III.** 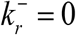; *α* = *β* = *χ*; *γ*_*r*_ = 0

In this case, **Eqs. 15** reduce to the following form which is exactly solvable.

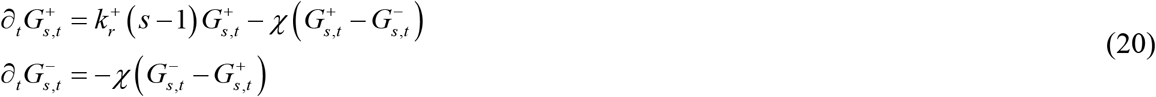

The general solution of **Eqs. 20** can be written as follows.

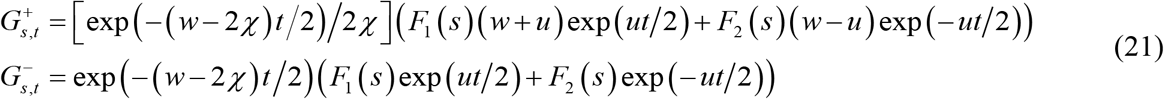

Here 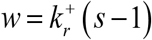 and 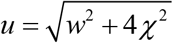. Noting the initial conditions 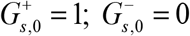 for a finite *χ* values one finds the following expressions for the functions *F*_1_(*s*) and *F*_2_(*s*).

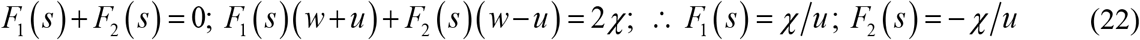

Upon using these, one can write down the particular solution of **Eqs. 20** as follows.

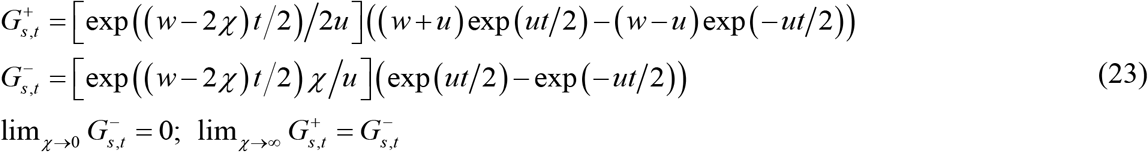

Using **Eqs. 22** and noting that 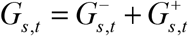 one can drive the following limiting conditions of the generating function.

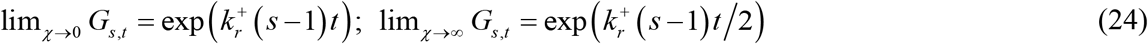

Using **Eqs. 23** one can directly derive the probability density function associated with the mRNA population at high and low values of the flipping rate as follows.

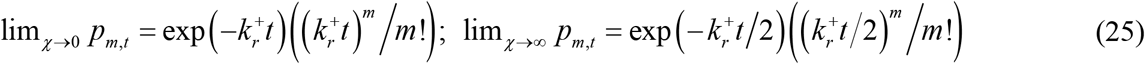

Further, one can derive the mean and the variance of the mRNA population explicitly as follows.

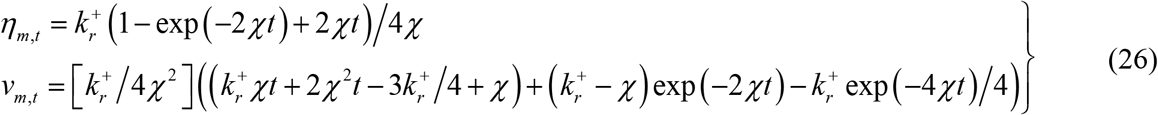

It is also interesting to note down the following limiting values.

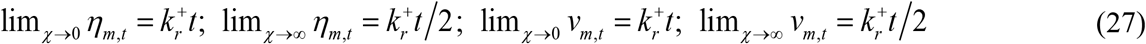

Remarkably, the functional form of the mRNA variance shows a turnover type behavior with respect to changes in *χ* and it has a definite maximum at the optimum flipping rate *χ*_*C*,*v*_. Explicit expression for this optimum *χ* can be derived from the following equation.

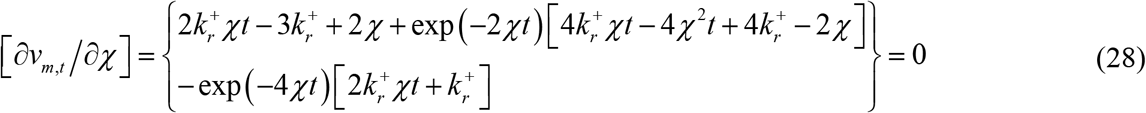

Upon ignoring the terms multiplying the exponentials for large values of *χ* and time as *t* → ∞, **Eq. 28** can be approximated to 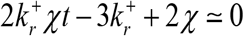 from which one finds the optimum value of *χ* at which the variance becomes a maximum as follows.

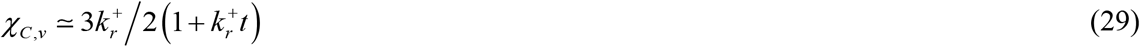

However, **Eq. 29** suggests that the optimum value of *χ* will be a time dependent quantity which decreases towards zero when the time increases towards infinity. Clearly, at the steady or stationary state i.e. in the limit as *t* → ∞, the existence of optimum flipping rate over the variance of the mRNA population will disappear. It is also interesting to note down the following limits.

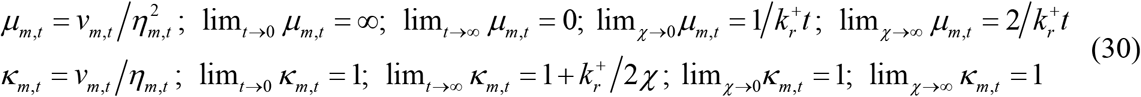

Remarkably, the Fano factor associated with the fluctuations in the number of mRNAs show a maximum deviation from the Poisson with respect to *χ* which can be demonstrated as follows.

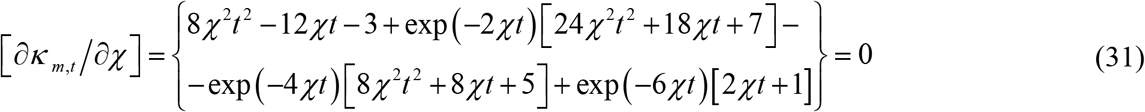

Similar to the derivation of **Eq. 29**, one can ignore the terms multiplying the exponentials and finally one obtains the following approximation for large values of *χ*.

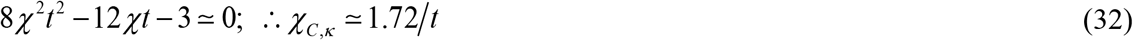

Clearly, at the steady state i.e. as *t* → ∞, the existence of optimum flipping rate over the Fano factor as well as the variance of the mRNA population will disappear. In other words, steady state theories and experiments cannot capture these important properties. Using detailed stochastic simulations, we will show in the later sections that both **Eqs. 29** and **32** are valid even when the conditions 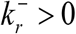; *α* ≠ *β*; *γ*_*r*_ > 0 are true. In these situations, we will show in the later sections that the optimum flipping rates (*α_C_*, *β_C_*) which maximize the variance and the Fano factor associated with the fluctuations in the number of mRNA molecules asymptotically approach their non-zero steady state limits.

### Dependency of the transcription rate on the on-off flipping dynamics

Let us consider a single transcription event. Here the length of the final mRNA transcript is *L* bp. Since the addition of each nucleotide at the ends of a growing mRNA is an energetically driven stochastic process, we can describe the entire transcription elongation as a directed walk with the microscopic transition rates 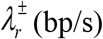. Here the superscript ‘+’ represents the on-state and ‘−’ represents the off-state of the promoter and 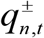 are probabilities of finding of the transcription machinery with *n* number of transcribed mRNA bases in the respective on-off state channels at time *t*. Now we can consider three possible scenarios as depicted in **Figs. 2A-B** viz. (a) when all the microscopic transitions of the transcription elongation event are characterized homogeneously with the rates 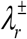 and there is no flipping across the on-off state channels, then one finds that 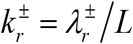 and subsequently the average resultant transcription rate will be equal to either 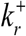 or 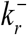 depending on the channel used for the transcription elongation (**Fig. 1**, **Figs. 2B1** and **2B3** respectively). On the other hand, when there is a flipping across on-off state channels, then the resultant transcription rate *ξ* will be somewhere in between 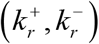 (**Fig. 2B2**). Clearly the transcription rate *ξ* associated with each mRNA transcript will be a random variable and its ensemble average across several individual transcripts is defined as *k_r_* = ⟨*ξ*⟩. When the timescales of flipping across on-off states are comparable with the timescales of the microscopic transcription elongation events, then such randomly interrupted transcription elongation can be well described by the following set of coupled master equations.

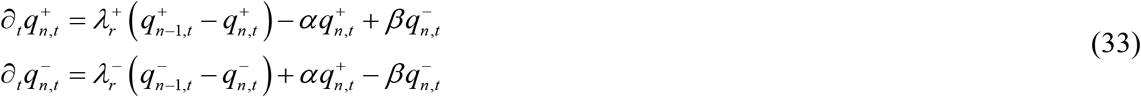

**FIGURE 2.**
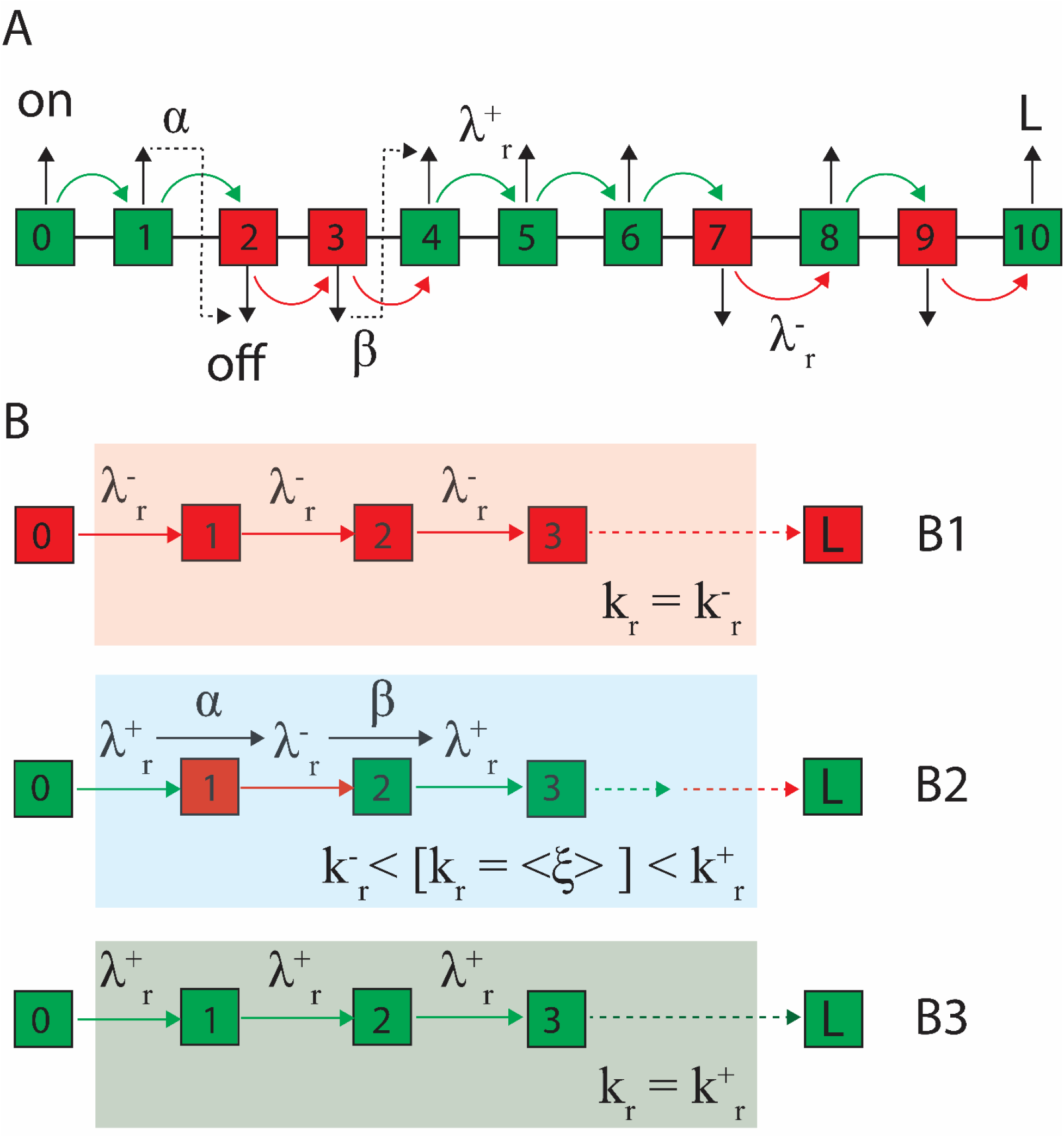
Stochasticity in the transcription rate. **A**. In this model the total length of the full transcript is *L* = 10 bp. Green color denotes the on-state and red color denotes the off-state channels. At time *t* = 0, the system was in the on-state. Green to green transition is characterized by the microscopic transcription elongation rate 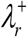 and red to red transition is characterized by the elongation rate 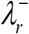. Here 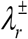 are measured in bp/s. **B**. When the entire transcription process follows the pure off-state channel (**B1**), then the overall transcription rate scales as 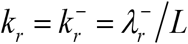 that is measured in *s*^−1^. When there is a flipping across the on- and off-state channels (**B2**), then the transcription rate of a given mRNA trajectory will be somewhere inside as 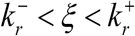. For a stall-continue type elongation of RNA polymerase, it approaches the limit 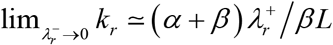 as shown in **Appendix A** where *k*_*r*_ = ⟨*ξ*⟩. When the entire transcription process follows the pure on-state channel (**B3**), then the overall transcription rate can be expressed as 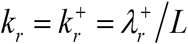.

The initial and boundary conditions associated with **Eqs. 34** can be given as follows [37–39].

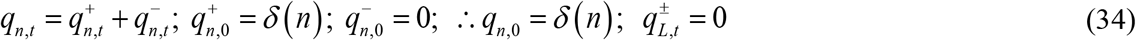

Upon defining 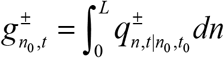 which is the overall probability of finding the system within *n* ∈ (0,*L*) at time *t* starting from *n* = 0, one can straightforwardly derive the following results.

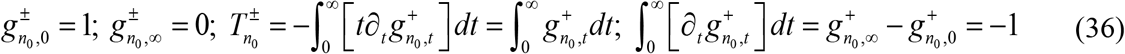

Further, we set that 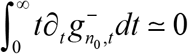 especially when 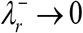 since the probability flux entering the off-state of the system will eventually return back to the on-state. The system can exit upon generating a complete transcript only from the on-state. Here 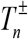 is the mean first passage times (MFPT) associated with the generation of a full mRNA transcript of length *L* starting from *n* number of initial bases of mRNA at time *t* = 0. One should note that the resultant transcription rate *ξ* is the inverse of the first passage time (FPT) which is a random variable. Clearly, the MFPTs 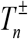 obey the following set of coupled backward type master equations [38, 39].

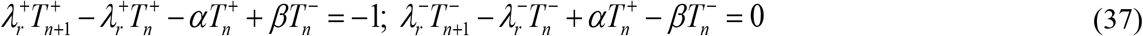

Here the boundary conditions directly follow from **Eqs. 34** as 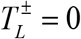. Upon solving **Eqs. 37**, the overall MFPT can be calculated as the sum 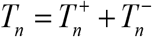. When the initial number of bases in the mRNA transcript is *n*_0_ = 0, then the required MFPT to generate a complete transcript starting from zero bases will be 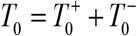 from which one can derive the expression for the overall average transcription rate as *k_r_* = [1 / *T*_0_]. The system of coupled **Eqs. 37** is exactly solvable. The solution corresponding to the first moment of the FPT 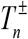 can be written as follows.

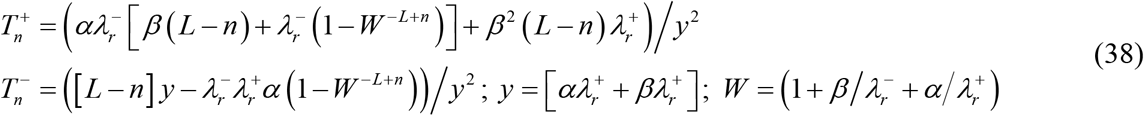

Here one should note the following limiting conditions.

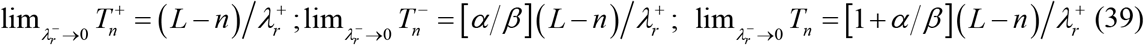

The second moment of the FPTs (we denote them as 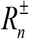) satisfy the following set of coupled backward type master equations [37].

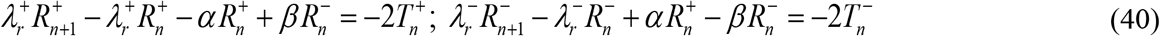

Here the boundary conditions of **Eqs. 40** directly follow from **Eqs. 34** as 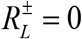. **Eqs. 40** can be derived using the following relationships.

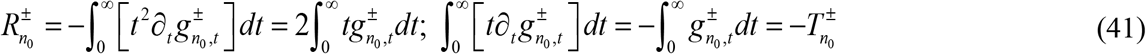

The set of coupled difference equations given in **Eqs. 40** along with **Eqs. 38** for the definition of 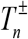 is exactly solvable and the complete solution is given in **Appendix A**. One can obtain the following simplified expressions for the second moments of FPTs associated with the generation of a complete mRNA transcript especially in the limit as 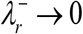.

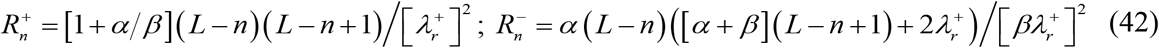

**Eqs. 42** can be obtained by substituting the expressions of 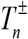 given by **Eqs. 39** in **Eqs. 40** subsequently solving the resultant difference equation. We denote the mean of FPTs with respect to *n* = 0 as 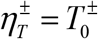 and, 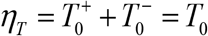 by definition. Using **Eqs. 39** and **42**, one can obtain the expressions for the variance (*v_T_*), Fano factor (*κ*_*T*_) and coefficient of variation (*μ*_*T*_) of the FPTs associated with the generation of a complete mRNA transcript of length *L* starting from *n* = 0 in the limit 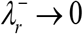 as follows.

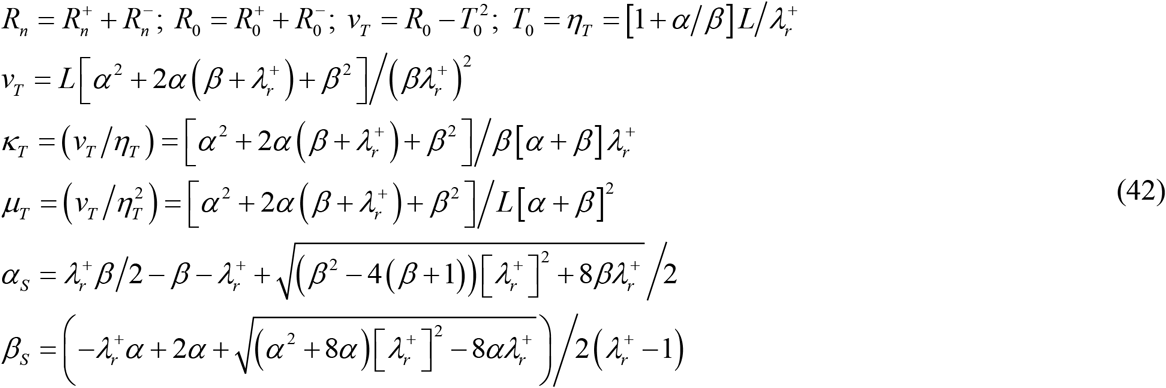

In **Eqs. 42**, (*α*_*s*_, *β*_*s*_) are the critical values of the flipping rates (which can be obtained by solving the equation *κ*_*T*_ = 1 for *α* by fixing *β* or for *β* by fixing *α*) such that when *α* < *α*_*S*_ or *β* < *β*_*S*_, then the Fano factor associated with the FPTs becomes as *κ*_*T*_ < 1 (sub-Poisson type). When *α* > *α*_*S*_ or *β* > *β*_*S*_, then *κ*_*T*_ > 1 (super-Poisson type). When *α* = *α*_*S*_ or *β* = *β*_*S*_, then *κ*_*T*_ = 1 (Poisson type). One should also note that *α*_*S*_ can exist only when 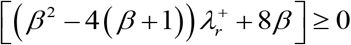 and *β*_*S*_ can exist only when 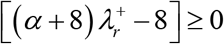. Since the overall transcription rate of a given mRNA transcript is the inverse of the respective FPT, one can conclude that the distribution of transcription rates will be the sub or super Poisson type depending on the relative values of the on-off flipping rates (*α*, *β*). Remarkably, the Fano factor *κ*_*T*_ associated with the distribution of FPTs is independent on the size of the mRNA transcript *L*. Further, the coefficient of variation of FPTs *μ*_*T*_ shows a turn over type behavior with respect to changes in *α* with a definite maximum at *α* = *β* which can be shown by solving *∂μ*_*T*_ / *∂α* = 0 for *α*. It is also interesting to note down the following limiting conditions.

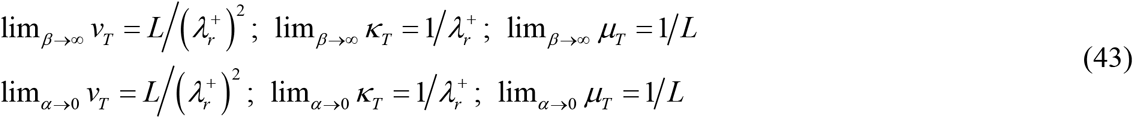

Especially, when *α* = *β* = *χ* then one can obtain the following simplified expression for *T_0_* using the set of **Eqs. 38** as follows.

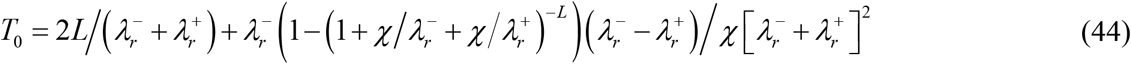

**Eqs. 44** clearly suggest that the overall average transcription rate *k_r_* is a function of the flipping rate *χ* especially when the timescale associated with the flipping dynamics is much slower than the timescale associated with the generation of a complete transcript. It is remarkable to note the limiting values of *T*_0_ as 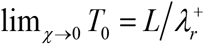 and, 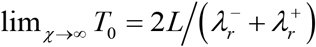. When 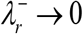 then **Eqs. 44** clearly suggest that *T_0_* will be independent of the on-off flipping rate *χ*. However, the MFPT will be doubled under such situations. When 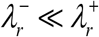 and for sufficiently large values of the transcript length *L*, one finally obtains the limiting conditions in line with **Eqs. 19** and **30** as 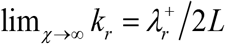 and, 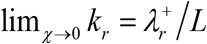 where 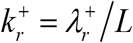. These limits follow directly from the initial conditions given in **Eqs. 35**. For example, when we hypothetically set up the initial conditions as 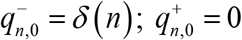, then we need to swap the on-off state superscript indices (+ and −) in **Eqs. 39** and subsequently one obtains the limiting values 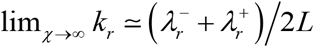 and, 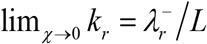 where 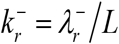 even when 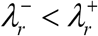.

When 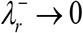 then the system of **Eqs. 37** will be uncoupled. In this situation, the overall average transcription rate *k_r_* rescales from 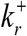 to 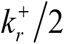 when *χ* tends towards infinity i.e. the overall transcription rate *k_r_* will be confined inside the interval 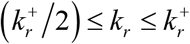. When 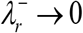 and *α* ≠ *β* then the overall average effective transcription rate can be expressed as the functions of the on-off flipping rates as 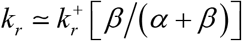. This equation suggests the probable range of *k_r_* as 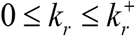 depending on the relative values of the on-off flipping rates. In the following sections, we investigate the possibility of various steady state solutions to **Eqs. 1**.

### Steady state solutions

Under steady state conditions, **Eqs. 1** reduce to the following form.

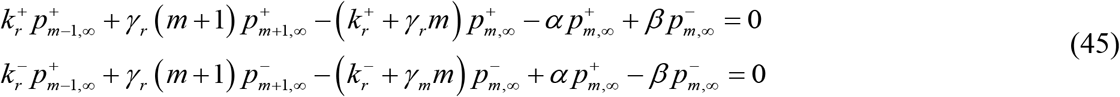

Here 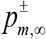 are the stationary state probabilities associated with the finding of *m* number of mRNAs in the respective on (+) and off (−) channels of transcription. **Eqs. 45** also can be solved using the standard generating function formalism. One can define the steady state generating functions and the corresponding probability densities as follows.

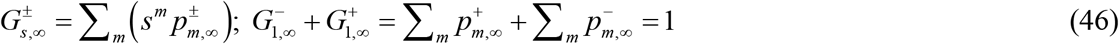

Using the generating functions defined in **Eqs. 46**, one can recover the respective probability density functions as follows.

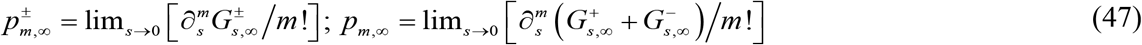

Upon applying the transformation given in **Eqs. 46** into **Eqs. 45**, one obtains the following set of coupled ordinary differential equations.

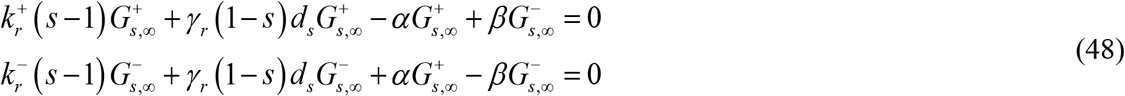

In Refs. [15, 17], the solutions to **Eqs. 45** were expressed in terms of confluent hypergeometric functions. Here we derive an alternate solution set that is easy to handle. **Eqs. 48** can be split into the following set of uncoupled ordinary differential equations.

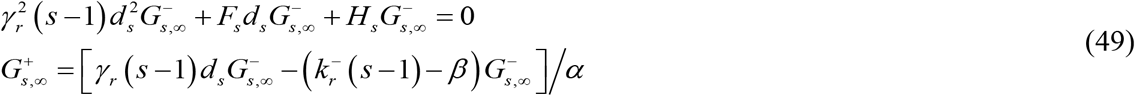

Various functions in Eqs. 49 are defined as follows.

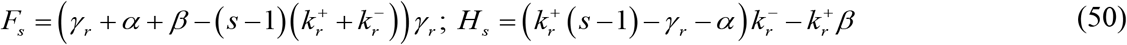

Using the substitution as 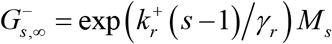, where *M_s_* is an arbitrary function of *s*, one can reduce **Eqs. 49** into the following form.

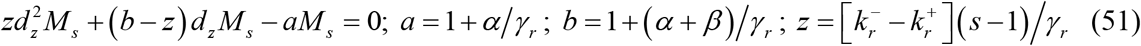

Solution to **Eq. 51** can be expressed in terms of Kummer functions as follows [40].

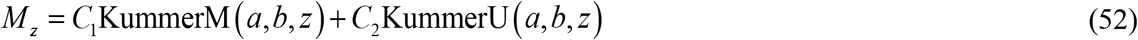

Since KummerU (*a*,*b*, 0) → ∞ for any arbitrary values of {*a*, *b*} > 0 in **Eq. 52**, we need to set *C*_2_ = 0 to enforce the normalization condition of the probability density function. We use the following properties of the KummerM functions to further simplify our results [40, 41].

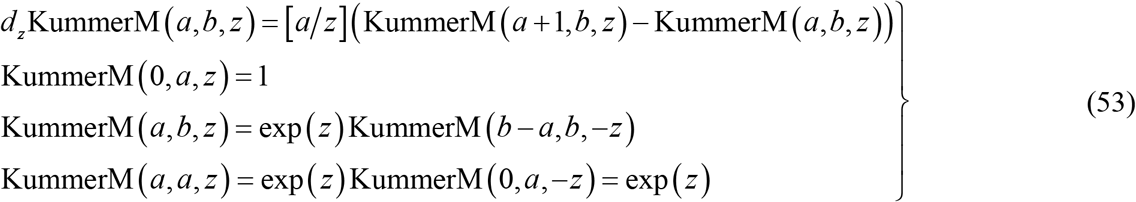

Using these properties, one finally obtains the solution to the generating functions obeying the set of differential equations **Eqs. 48** as follows.

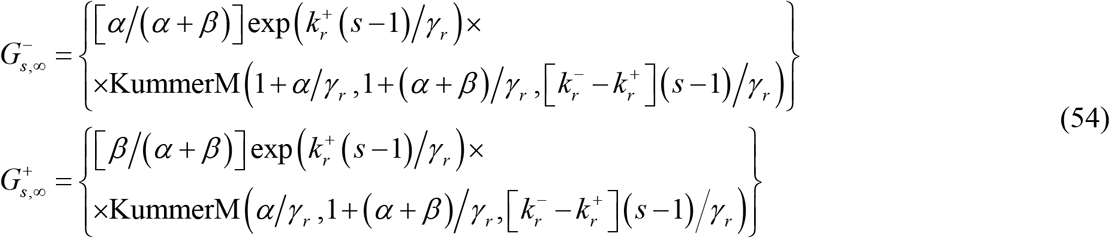

Here the KummerM function can be defined explicitly as follows [40, 41].

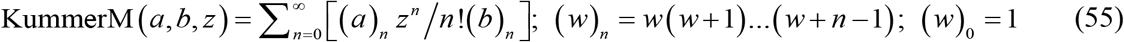

Upon expanding the generating functions in terms of Macularin series with respect to *s* around *s* = 0 and then substituting *s* = 1 in the computed series, one finally obtains the expressions for the respective probability density functions as follows.

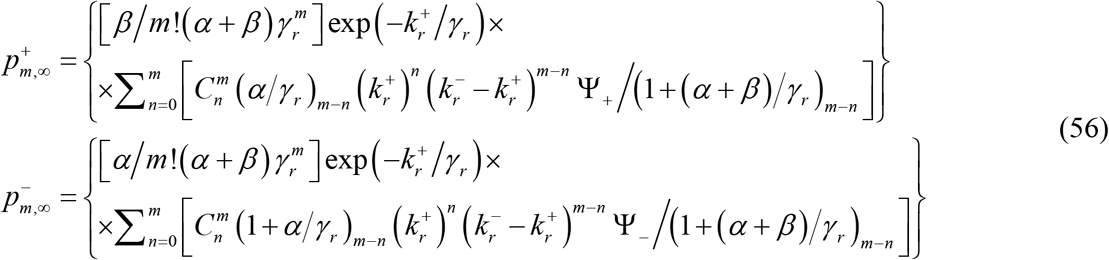

Various paramaters and terms in **Eqs. 56** are defined as follows.

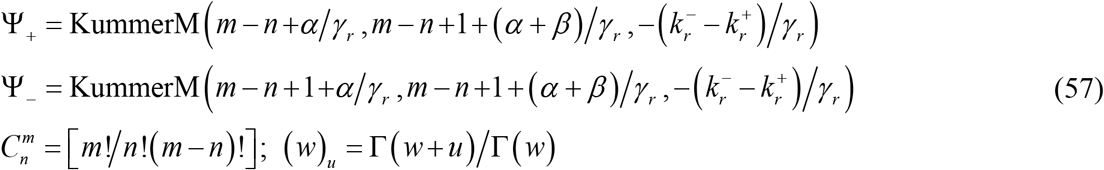

From **Eqs. 54**, one can derive the expressions for the steady state mean, variance, coefficient of variation and Fano factor (we denote them as *η*_*m*,∞_, *v*_*m*,∞_, *μ*_*m*,∞_, *κ*_*m*,∞_ respectively) as follows.

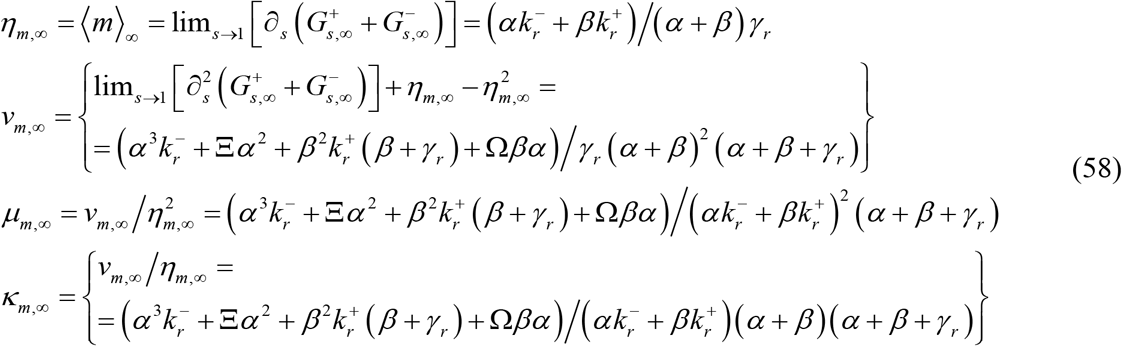

Various terms in **Eqs. 58** are defined as follows.

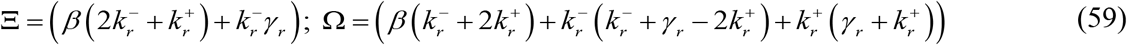

**Eqs. 58** suggest that, in the presence of flipping across the on-off states, the effective transcription rate transforms as 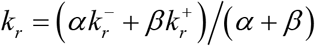 under steady state conditions. In the following sections we will consider various cases of approximations to **Eqs. 54**-**59**.

**Case I**. *α* ≠ *β*; 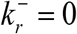

In this situation, **Eqs. 54** reduce to the following form.

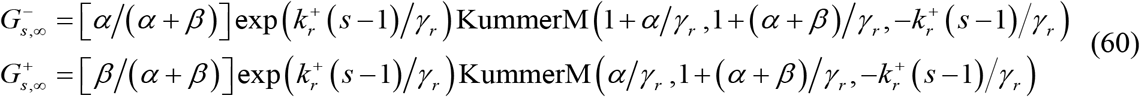

Upon expanding these generating functions in terms of Macularin series around the point *s* = 0 and then substituting *s* = 1 in the computed series, one finally obtains the following probability density functions associated with the steady state mRNA populations.

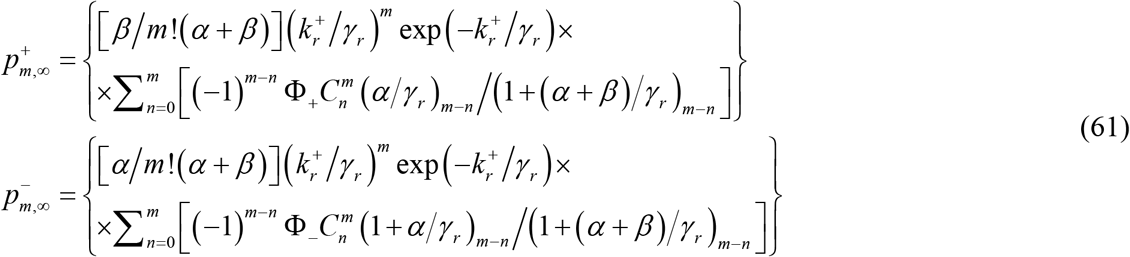

Various terms in **Eqs. 61** are defined as follows.

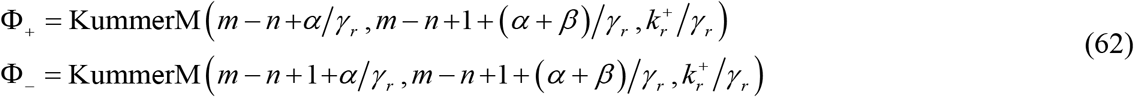

From **Eqs. 60**, one can derive the expressions for the steady state mean, variance, coefficient of variation and Fano factor as follows.

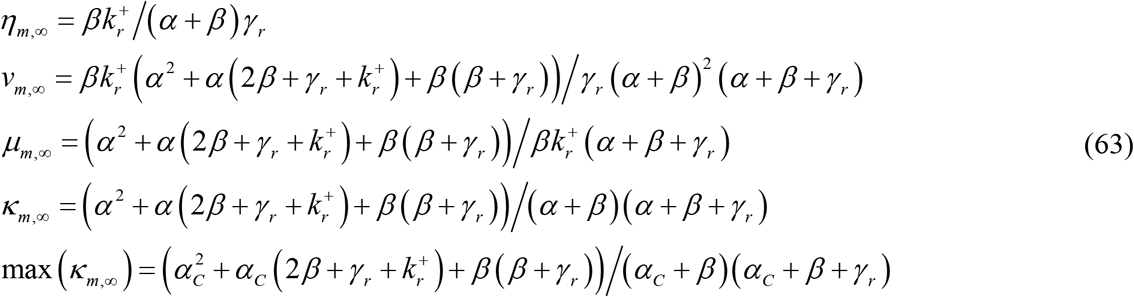

Clearly, when 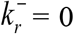, then the overall transcription rate rescales as 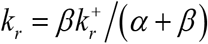 and using the respective limiting condition from **Eqs. 63** one can derive the optimum *α* at which the stationary state Fano factor *κ*_*m*,∞_ attains a maximum as 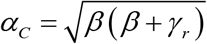. This can be obtained by solving *∂κ*_*m*,∞_ / *∂α* = 0 for *α*. Using the definition of the burst size or the transcription efficiency *σ* = *k_r_* / *α*, one can also show that the steady state Fano factor *κ*_*m*,∞_ will attain a maximum value at the optimum transcription efficiency 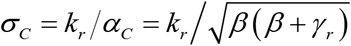.

**Case II**. *α* = *β* = *χ*; 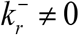

In this case, the generating functions reduce to the following form.

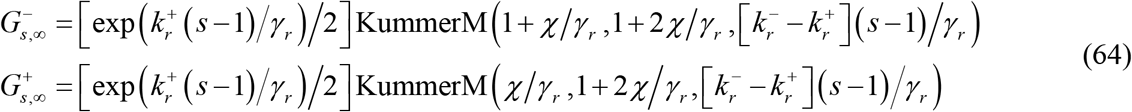

Upon expanding these generating functions in terms of Macularin series around the point *s* = 0 and then substituting *s* = 1 in the computed series, one finally obtains the following probability density functions associated with the steady state mRNA populations.

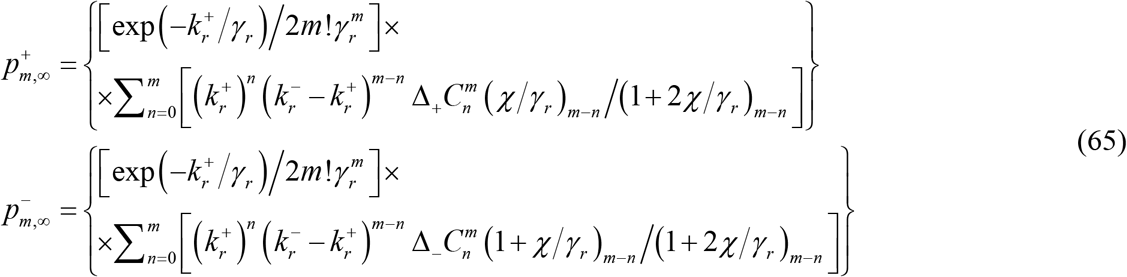

Various terms and functions in **Eqs. 65** are defined as follows.

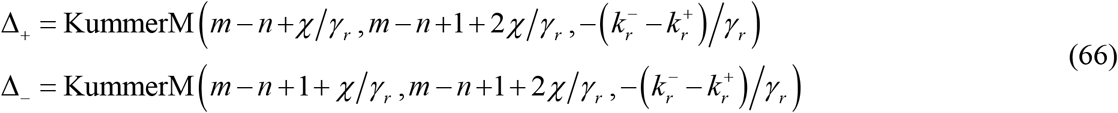

Using the generating functions given in **Eqs. 64** and noting the definition of the overall generating function 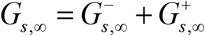, one can derive the following properties of the stationary state mRNA populations and their various limiting properties.

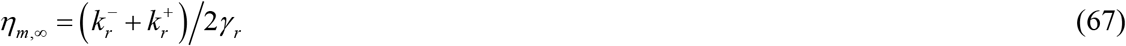

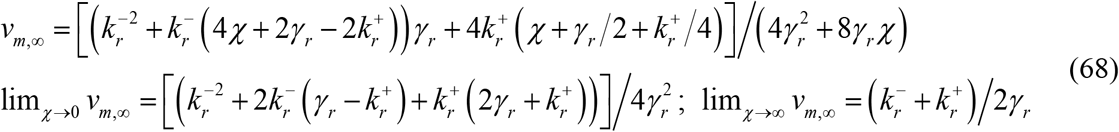

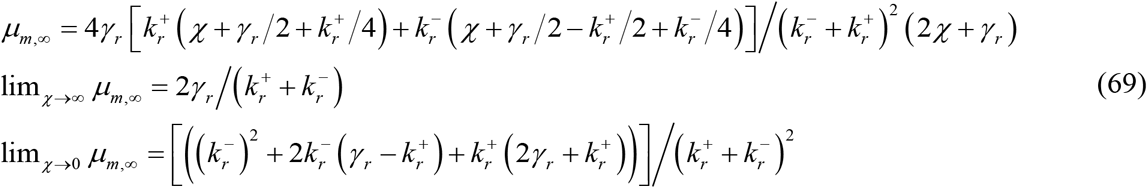

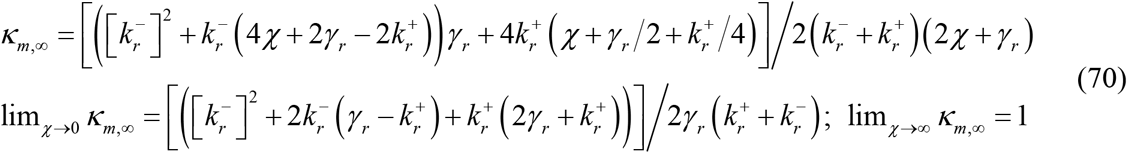

**Case III**. *α* = *β* = *χ*; 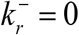

In this situation, various statistical properties of the stationary state mRNA populations and their various limiting values can be derived as follows.

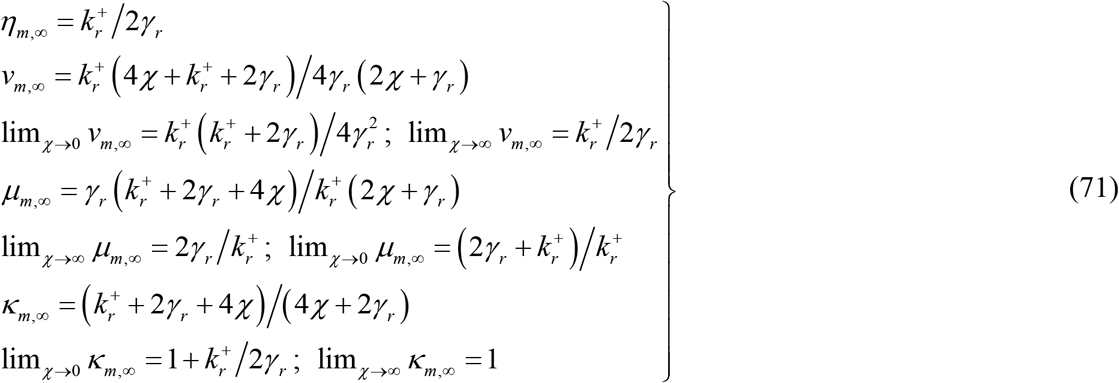

**Case IV**. *α* ≠ *β*; *α* ≪ *γ*_*r*_; *β* ≪ *γ*_*r*_; 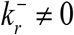

In this case, the generating functions reduce to the following simple form.

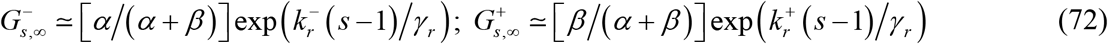

The corresponding probability density functions can be derived via expanding these generating functions in terms of Macularin series around *s* = 0 and then substituting *s* = 1 in the computed series expansion.

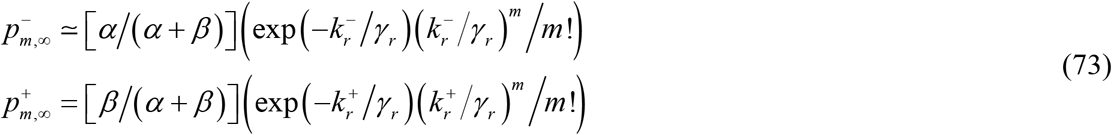

Therefore, one finally obtains the following bimodal Poisson type expression for the probability density function associated with the stationary state mRNA populations.

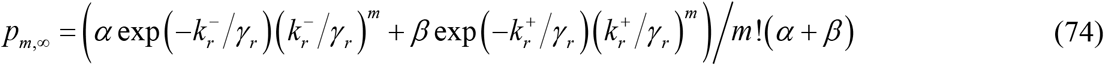

These types of bimodal probability density of the mRNA populations seem to play critical role in the cell to cell variability within an organism [14]. From **Eqs. 60**, one can derive the expressions for the steady state statistical properties of the mRNA population as follows.

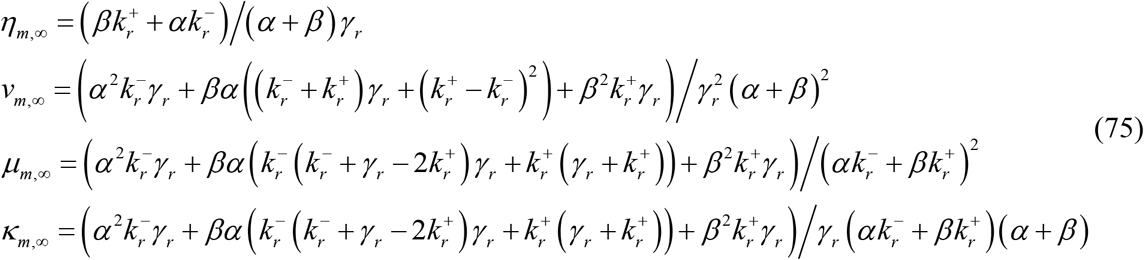

From **Eqs. 75** one can conclude that the steady state Fano factor will attain a maximum when the at the flipping rates *α_C_*, *β_C_* which are connected via 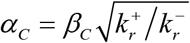.

**Case V**. *α* ≠ *β*;; *α* ≪ *γ*_*r*_; *β* ≪ *γ*_*r*_; 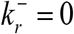

This is exactly the scenario addressed in Ref. [14]. In this case, the generating functions defined in **Eqs. 54** takes the following simple form.

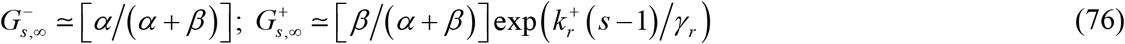

We have used the properties of the KummerM functions defined in **Eqs. 53** to derive **Eqs. 76**. The corresponding probability density functions can be derived as follows.

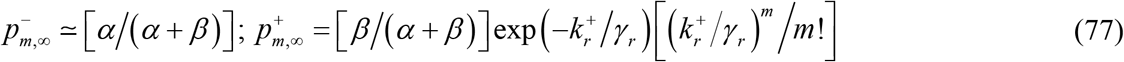

Using **Eqs. 77**, one finally obtains the Poisson density function with zero spike [14].

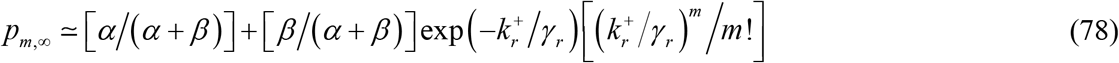

In **Eq. 78**, the number of mRNA molecules *m* takes only the integer values and **Eq. 78** is valid for the entire range of *m* i.e. *m* = 0 to infinity. Noting that 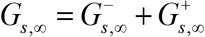, from **Eqs. 76**, one can derive various statistical properties such as mean, variance, coefficient of variation and Fano factor associated with the stationary state mRNA populations as follows.

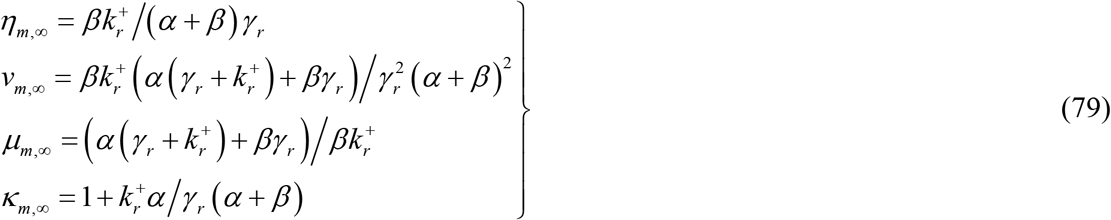

## STOCHASTIC SIMULATION METHODS

To check the validity **Eqs. 29** and **32** under various conditions, we performed detailed stochastic simulations on the complete system of master equations **Eqs. 1**. Here there are two different timescales viz. the timescale associated with the generation of a complete mRNA transcript and the timescale associated with the flipping dynamics across on-off state channels of transcription. We used the Gillespie algorithm [42, 43] to simulate the system of **Eqs. 1**. Let us denote the number of mRNA molecules at time *t* as *m*. Initially at *t* = 0, *m* = 0 and the system was in the on-state so that 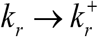. Clearly, there are four different reaction transitions viz. (*m* − 1) → *m* which represents the zeroth order transcription with a rate *k_r_*, (*m* + 1) → *m* which represents the first order recycling of mRNA molecules with a rate *γ*_*r*_ *m*, [+] → [−] represents the flipping from the on-state to the off-state with a rate *α* and [−] → [+] represents the flipping from the off-state to the on-state with a rate *β*. The total reaction flux here is *f_T_* = *k_r_* + *γ*_*r*_ *m* + *α* + *β*. We generated two different random numbers which are equally distributed inside *r*_1_, *r*_2_ ∈(0,1). The reaction times were sampled from the exponential type distribution *p* (*τ*) ∝ exp(− *f*_*T*_*τ*). This can be achieved via transforming *r_1_* using the rule *τ* = − (ln *r*_1_)/*f*_*T*_. We used *r_2_* to decide on which reaction takes place at this time point. In these iteration steps we set 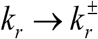 depending on the current state of the transcription channel.

a. When *r*_2_ ≤ (*k_r_*/ *f_T_*) then the transcription reaction will take place.
b. When (*k_r_* / *f_T_*) < *r*_2_ ≤ ((*k_r_* + *γ*_*r*_*m*)/*f*_*T*_), then the recycling of mRNA will take place.
c. When ((*k_r_* + *γ*_*r*_*m*)/*f_T_*) < *r*_2_ ≤ ((*k_r_* + *γ*_*r*_*m* + *α*)/ *f_T_*), then the transition from the on-state to the off-state will take place. Upon such transition we set the transcription rate as 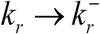.
d. When ((*k_r_* + *γ*_*r*_*m* + *α*)/ *f_T_*) < *r*_2_, then the transition from the off-state to the on-state will take place. Upon such transition we set the transcription rate as 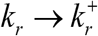.

Several trajectories were generated each with a total time *t_T_* and various statistical properties of mRNA numbers such as mean, variance, coefficient of variation and Fano factor were computed across the time axis. To compute the optimum flipping rate, same set of simulations were performed at different values of the flipping rate. Using this dataset, the optimum flipping rates at which the variance and the Fano factor attained the maximum were numerically computed.

To compute the mean first passage time (MFPT) associated with the generation of a complete mRNA transcript we used the following algorithm. We set up the initial number of mRNA bases as *n* = 0 and the system starts from the on-state or off-state channel with the respective microscopic elongation rates 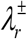. The completely transcribed mRNA transcript will have *L* number of bases. Clearly, there are three different transitions viz. (*n* − 1) → *n* which represents the transition with rate 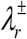 depending on the transcription channel at that time point, [+] → [−] represents the flipping from the on-state to the off-state with a rate *α* and [−] → [+] represents the flipping from the off-state to the on-state channel with a rate *β*. The total reaction flux here is *f_Q_* = *λ*_*r*_ + *α* + *β* where 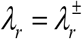 depending on the current state of the transcription channel. We generated two different random numbers which are equally distributed inside *r*_1_, *r*_2_ ∈(0,1). The reaction times were sampled from the exponential distribution *p* (*τ*) ∝ exp(− *f*_*Q*_*τ*). This can be achieved via transforming *r_1_* using the rule *τ* = −(ln *r*_1_)/*f_Q_*. We used *r_2_* to decide on which reaction transition takes place at this time point. In these iteration steps, we set 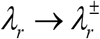 depending on the current state of the transcription channel.

a. When *r*_2_ ≤ (*λ*_*r*_/*f_Q_*) then the microscopic elongation transition *n* → (*n* +1) will take place.
b. When (*λ*_*r*_/*f_Q_*) < *r*_2_ ≤ ((*λ*_*r*_ + *α*)/*f_Q_*), then the transition from the on-state to the off-state will take place. Upon such transition we set the microscopic elongation transition rate as 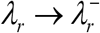.
c. When ((*λ*_*r*_ + *α*)/*f_Q_*) < *r*_2_, then the transition from the off-state to the on-state will take place. Upon such transition we set the microscopic elongation transition rate as 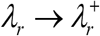.

When *n* = *L*, then the iteration was stopped and the first passage time (FPT) was noted. Several such trajectories were generated and the obtained FPTs were used to compute the MFPT and other statistical properties of FPTs such as variance, coefficient of variation and the Fano factor. This analysis was repeated at various of values of the flipping rates.

### RESULTS AND DISCUSSION

Transcription bursting seems to emerge as a consequence of the interplay between the on-off flipping rates (*α*, *β*), microscopic transcription elongation rates 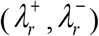, resultant transcription rates 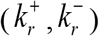, and the recycling rate of mRNAs *γ_r_*. When the on-off flipping rates and the transcription elongation rates are comparable to each other, then a continuous type transcription with monomodal type distribution of the mRNA populations will be observed as in **Fig. 3A**. In this case, the transcription will be always in the on-state by definition. Transcription bursting emerges when one sets *α* > *β* and the condition 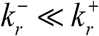 is satisfied which is clearly demonstrated in **Figs. 3B-D**. Here *β* is the rate of flipping from the off-state to the on-state of transcription and *α* is the rate of flipping from the on-state to the off-state. In most of the natural scenarios one finds that 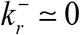. Therefore, the inequality condition *α* > *β* along with the decay rate constant *γ_r_* decides the emergence of the bursting in transcription and also the burst size that is defined as *σ* = *k_r_*/ *α*. The mRNA number fluctuations in the two-stage or continuous transcription and decay follows a typical Poisson density function with Fano factor = 1. When the transcription undergoes on-off flipping dynamics, then the mRNA number fluctuations follow a typical super Poisson type density function with a Fano factor more than one. Earlier studies suggested that such systems are closely follow a negative binomial distribution function [15]. The entire transcription scheme can be fragmented into several sub-processes viz.

a. The flipping dynamics across the on-off channels of the transcription. This is characterized by the flipping rates (*α*, *β*). Here the on-state describes the fully functional transcription machinery and the off-state describes a scenario where the RNAP is stalled due to the presence huge positive supercoil barrier ahead or other chromosomal barriers.
b. Microscopic transcription elongation events representing the growth of individual mRNAs starting from *n* = 0 number of bases toward *n* = *L* bp. This is characterized by the microscopic transcription elongation rates 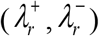 corresponding to the on and off channels respectively.
c. Mesoscopic transcription dynamics. This is characterized by the mesoscopic transcription rates 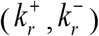 corresponding to the on and off state channels which are the cumulative effects of (**a**) and (**b**). When the transcription follows pure on or off state channels so that there is no flipping across them i.e., *α* = *β =* 0, then one finds that 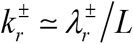. When the transcription follows a random trajectory via flipping across on and off states, then the resultant transcription rate *ξ* will be a random variable. Clearly, *ξ* varies across different mRNA transcripts. The overall average transcription rate can be defined as *k_r_* = ⟨*ξ*⟩. When *α* ≠ *β,* then one finds the probable range of the resultant transcription rate 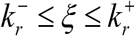 and 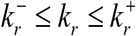.
d. Recycling dynamics of mRNAs which is characterized by the decay rate *γ_r_*. Independently it is a first order decay which follows a Poisson type density function.

**FIGURE 3.**
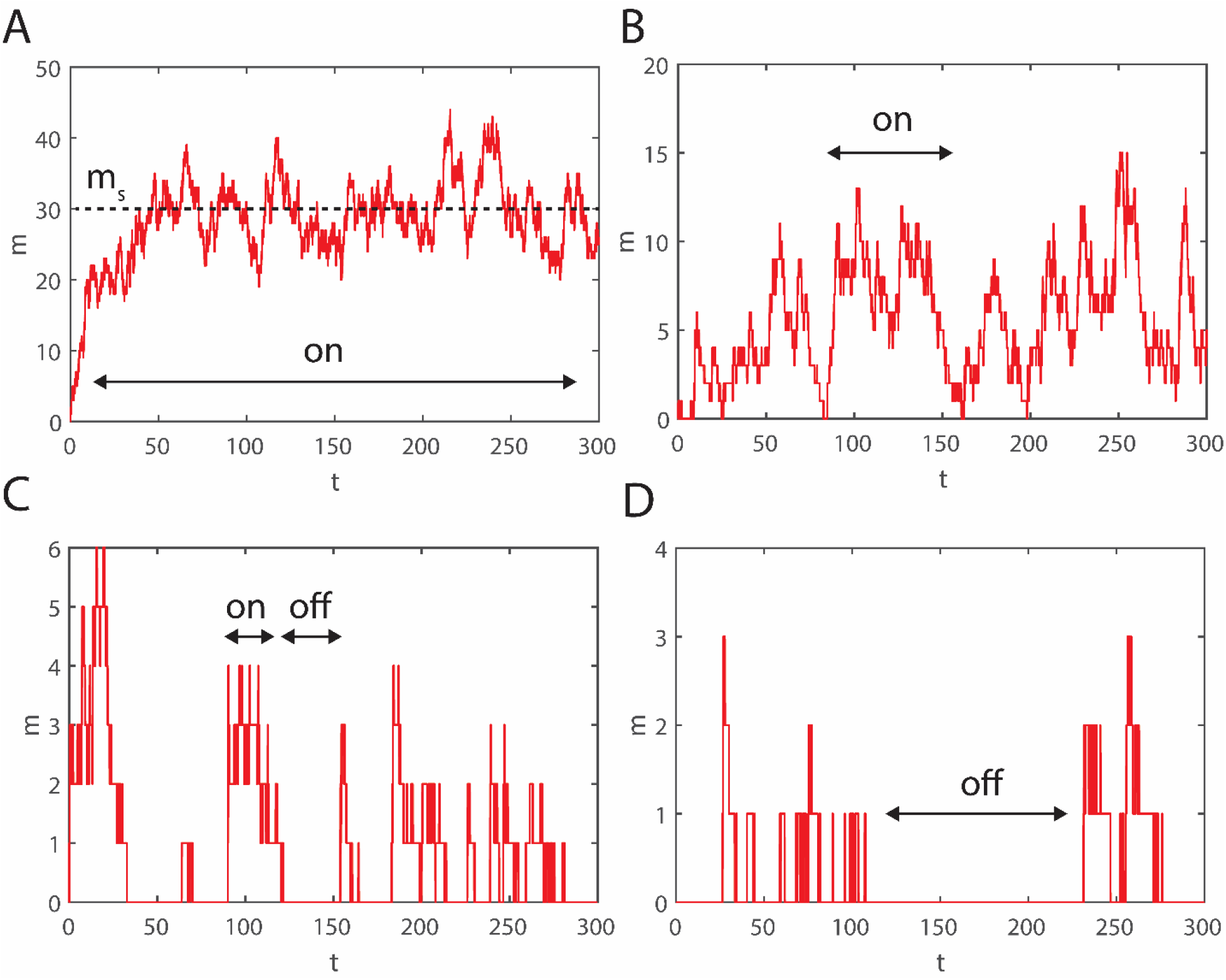
Emergence of the transcription bursting phenomenon. These are all single stochastic trajectories. **A**. Continuous transcription process. Here the parameters settings are *α* = *β* = 10^−6^ s^−1^, 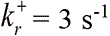, *γ*_*r*_ = 0.1 s^−1^, and 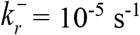. In these settings, one finds that 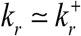 and therefore the steady state mRNA numbers will be 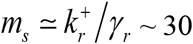. Initially the system was set into the on-state of transcription. **B**. Transcription bursting emerges when one sets high value for *α* and low value for *β* apart from the mandatory condition that 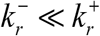. Here *α* = 3, *β* = 1 and rest of the parameters are set as in in the panel **A**. In **C**. *β* = 0.1 and in **D**. *β* = 0.05 and rest of the other parameters are set as in the panel **A**.

Individually, the population of states in the on-off channels associated with the uncoupled sub-process (**a**) follows a binomial density function for which the Fano factor is less than one. This is similar to the tossing of a coin. On the other hand, the population of various states in the sub-processes (**b**), (**c**) and (**d**) individually follow typical Poisson density function with a Fano factor of one (see **Appendix B** for a simplified derivation). When the sub-process (**a**) is dynamically coupled with (**b**) and (**c**), then the resultant or effective transcription rate *ξ* randomly fluctuates among the population of transcription events across 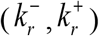. The probability density function associated with the fluctuations in the resultant transcription rate *ξ* will be dependent on the on-off flipping rates, microscopic elongation rates 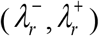 and the length of the mRNA transcript *L*.

**Figs. 4** and **5** demonstrate how the statistical properties of the mRNA number fluctuations varies with time and flipping rate parameters. When *α* = *β* = *χ* and *γ_r_* = 0, then one finds from **Figs. 4B** and **D** that the variance and the Fano factor associated with the mRNA numbers attain maxima at the optimum flipping rates *χ*_*C, v*_ and *χ*_*C,κ*_ respectively. These optimum flipping rates seems to be decrease with time towards zero in line with the theoretical predictions of **Eqs. 29** and **32**. These results are shown in **Figs. 5**. Clearly, the system will follow a Poisson type density function (Fano factor of mRNA number fluctuations equals to one) when the flipping rates become *χ* → 0 as well as *χ* → ∞. This reasonable since when *χ* → 0 then the on-off channels will be uncoupled and all the transcription events will be via the on-state channel. When *χ* → ∞ then the system of strongly coupled and both the transcription channels will be equally utilized. When *γ_r_* ≠ 0, then the optimum flipping rates *χ*_*C,v*_ and *χ*_*C,κ*_ first converge to the steady state limits and then move slowly towards zero which is evident from **Figs. 4E, G**, **Figs. 5B** and **E**. **Figs. 6** demonstrate how the statistical properties of the mRNA number fluctuations vary with time when *α* ≠ *β*. Remarkably, **Eqs. 63** suggested that there exists steady state optimum points *α_C_* and *β_C_* at which the Fano factor attains the maximum. These optimum points are connected via 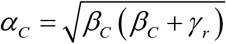 as shown in **Eqs. 63** and **Figs. 7**. When we fix *α* and iterate *β*, then one finds the optimum value of *α* at which the Fano factor attains maximum as 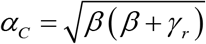. Particularly for a fixed *β* = 1, one finds for *γ_r_* = 1 that 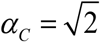 as shown in **Fig. 6D**. In the same way, when we fix *β* and then iterate *α*, then one finds the optimum *β* at which the Fano factor attains a maximum as 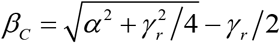 and so on. Particularly for a fixed *α* = 1, one finds for *γ_r_* = 1 that *β*_*C*_ = 0.618 as shown in **Fig. 6H**.

**FIGURE 4.**
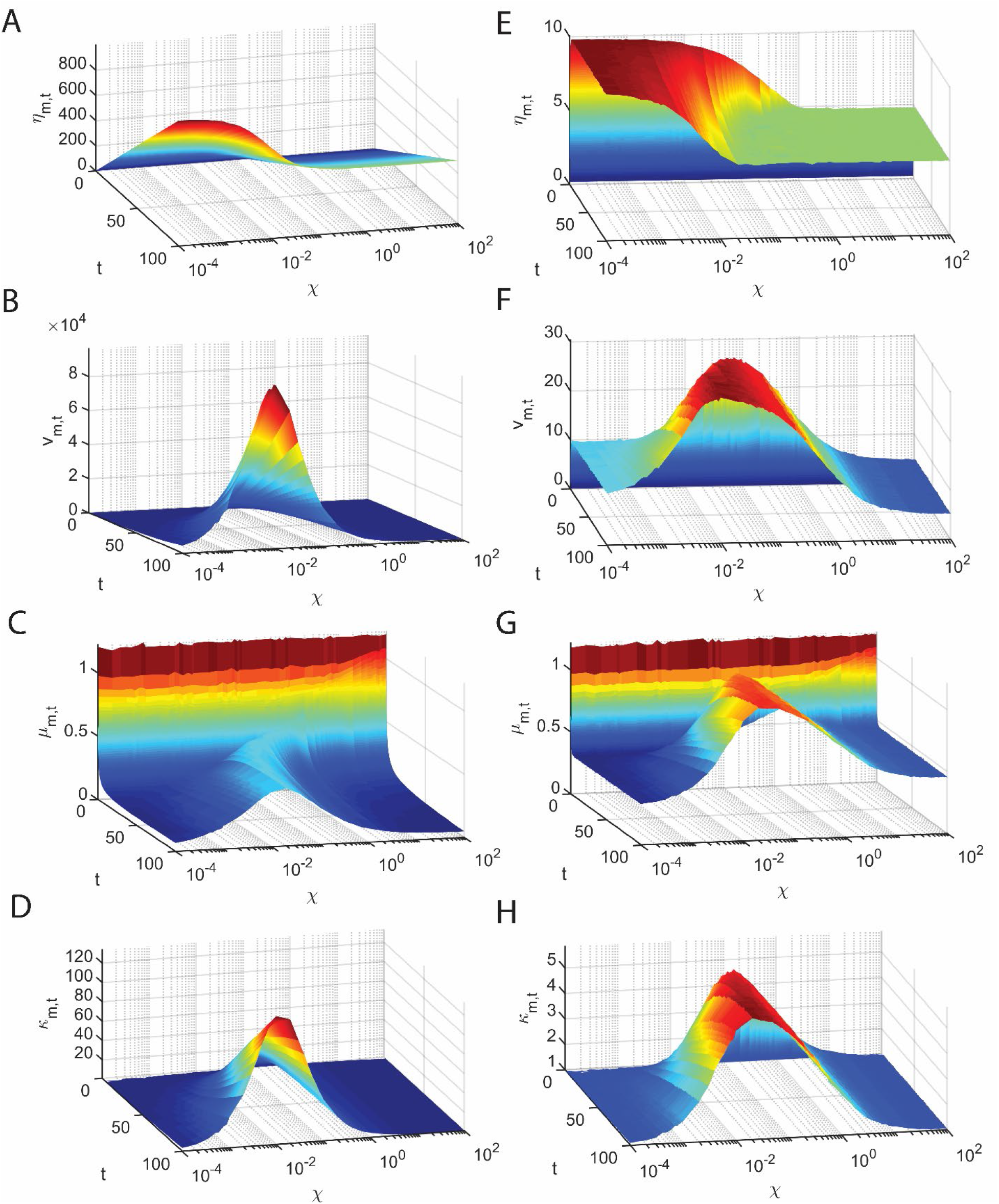
Simulation results on various time dependent statistical properties of mRNA number fluctuations in the pre-steady state regime of the transcription event. Here we have set *α* = *β* = *χ* for simplification, *η*_*m,t*_ denotes the mean, *κ*_*m,t*_ denotes the Fano factor, *μ*_*m,t*_ denotes the coefficient of variation, and *v*_*m,t*_ denotes the variance. In the panels **A-D** the parameter settings are 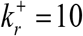, 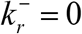 and *γ*_*r*_ = 0. In line with the predictions of **Eqs. 29** and **32** one observes maxima in the Fano factor and the variance with respect to the flipping rate *χ* which is also a time dependent quantity. As predicted by these equations, the optimum flipping rate shifts towards zero as time tends towards infinity. In the panels **E-H** the parameter settings are 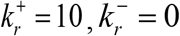 and *γ*_*r*_ = 1. Statistical properties were computed over 10^6^ number of trajectories.

**FIGURE 5.**
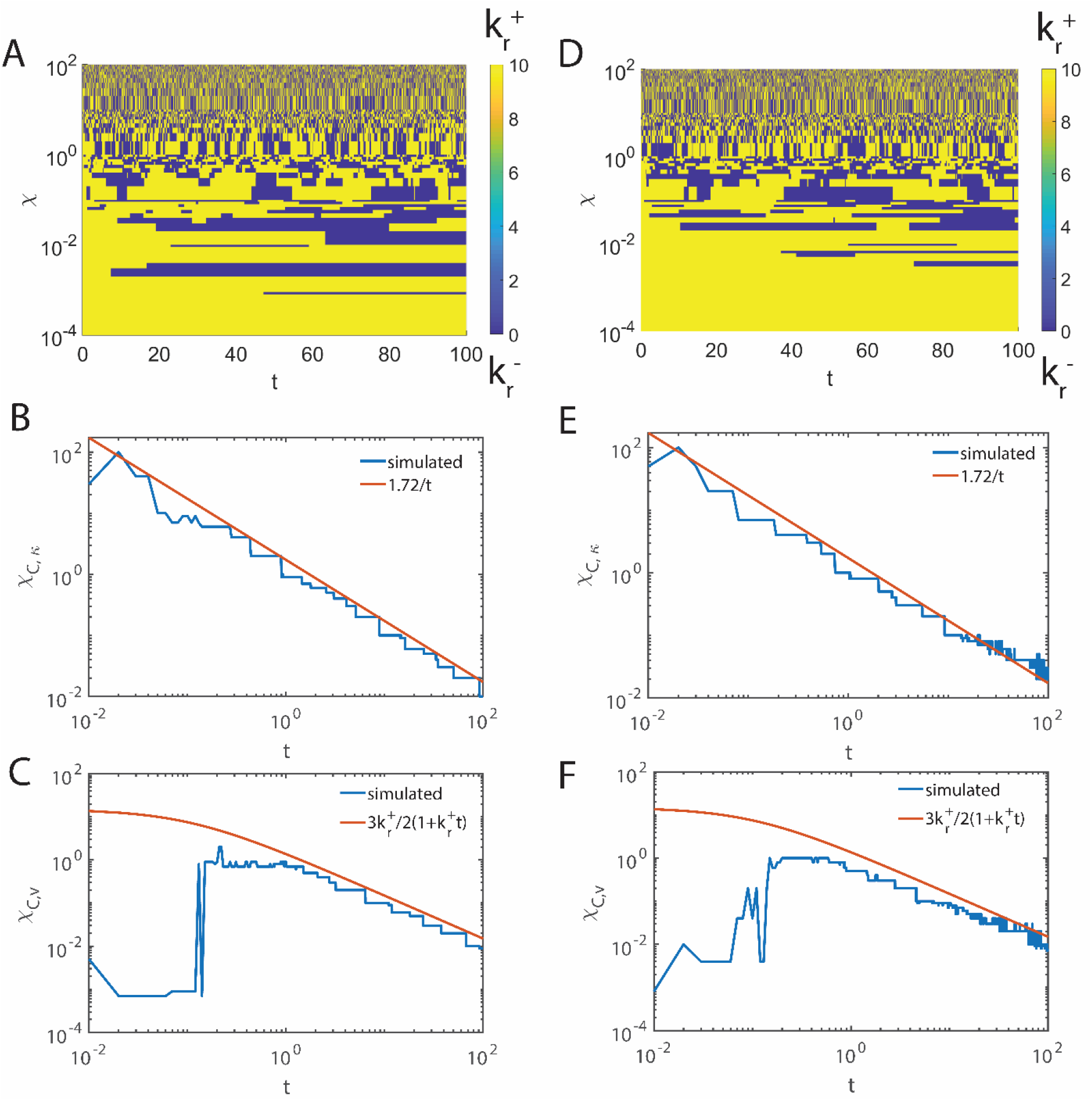
Variation of the optimum flipping rate *χ* that maximizes the variance and the Fano factor of mRNA numbers. In panels **A-C** the settings are 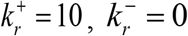 and *γ*_*r*_ = 0. In panels **D-F** the settings are 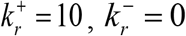 and *γ*_*r*_ = 1. Panels **A** and **D** represent the flipping dynamics across the on and off states. Panels **B** and **E** represent the variation of the optimum flipping rate with respect to time which maximize the Fano factor associated with the mRNA number fluctuations. Panels **C** and **F** represents the variation of the optimum flipping rate with respect to time which maximizes the variance associated with the mRNA number fluctuations. Here the solid red lines are the predictions by **Eqs. 29** and **32** which are valid when *α* = *β* = *χ*, 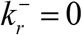 and *γ*_*r*_ = 0. In panels **C** and **F**, the precision in numerically computing the maximum points is lost when *t* < 0.1 since the surface is almost flat. Statistical properties were computed over 10^6^ number of trajectories.

**FIGURE 6.**
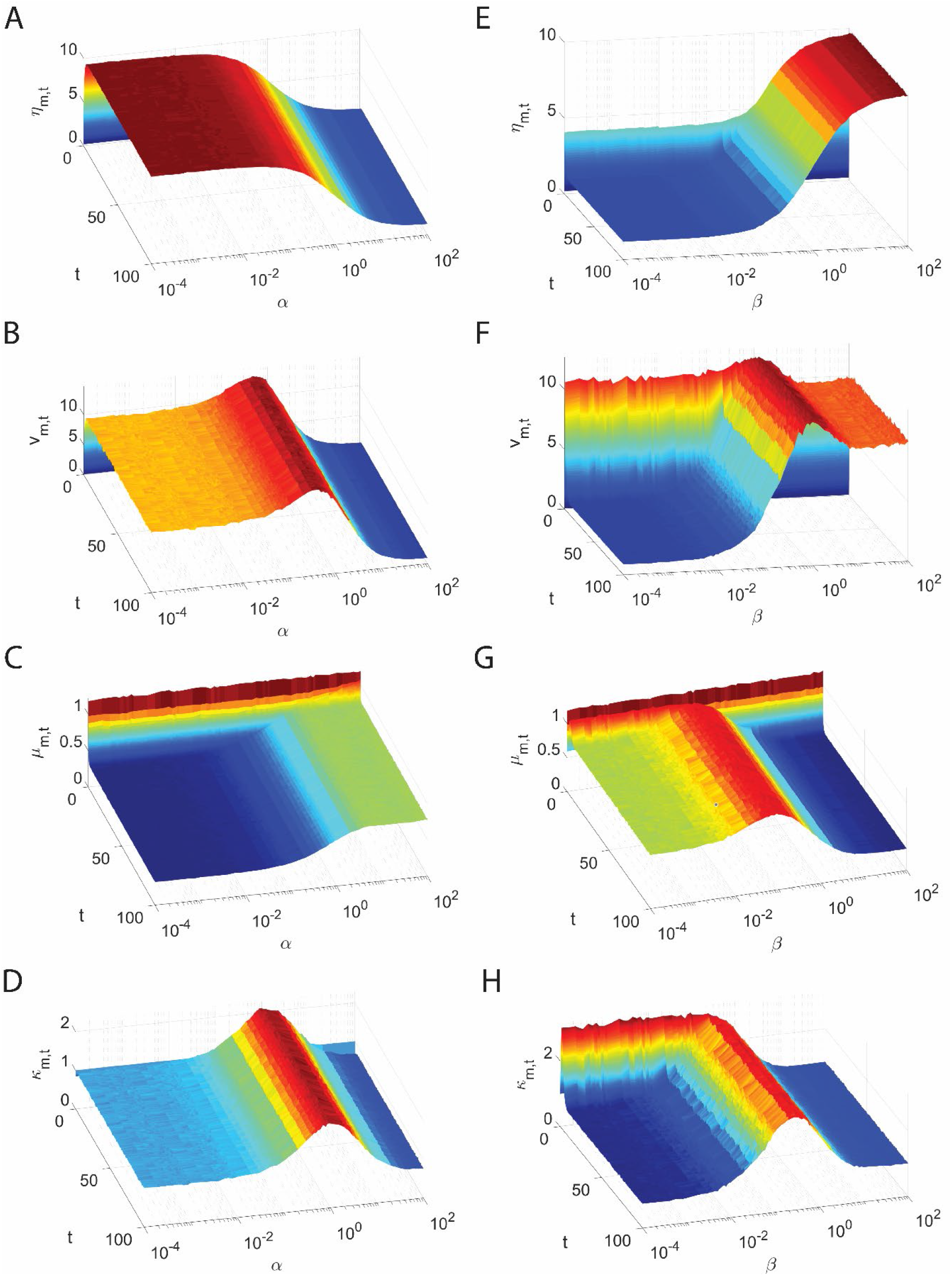
Various time dependent statistical properties of mRNA fluctuations in the pre-steady state regime of the transcription event. Here we have set *α* ≠ *β*, *η*_*m,t*_ denotes the mean, *κ*_*m,t*_ denotes the Fano factor, *μ*_*m,t*_ denotes the coefficient of variation, and *v*_*m,t*_ denotes the variance of the mRNA number fluctuations. In the panels **A-D** the settings are 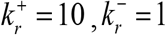, *γ*_*r*_ = 1 and *β* = 1. In the panels **E-H** the settings are 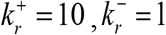, *γ*_*r*_ = 1 and *α* = 1. In line with the predictions of **Eqs. 29** and **32** one observes maxima in the Fano factor and the variance even when *α* ≠ *β*. Statistical properties were computed over 10^6^ number of trajectories.

**FIGURE 7.**
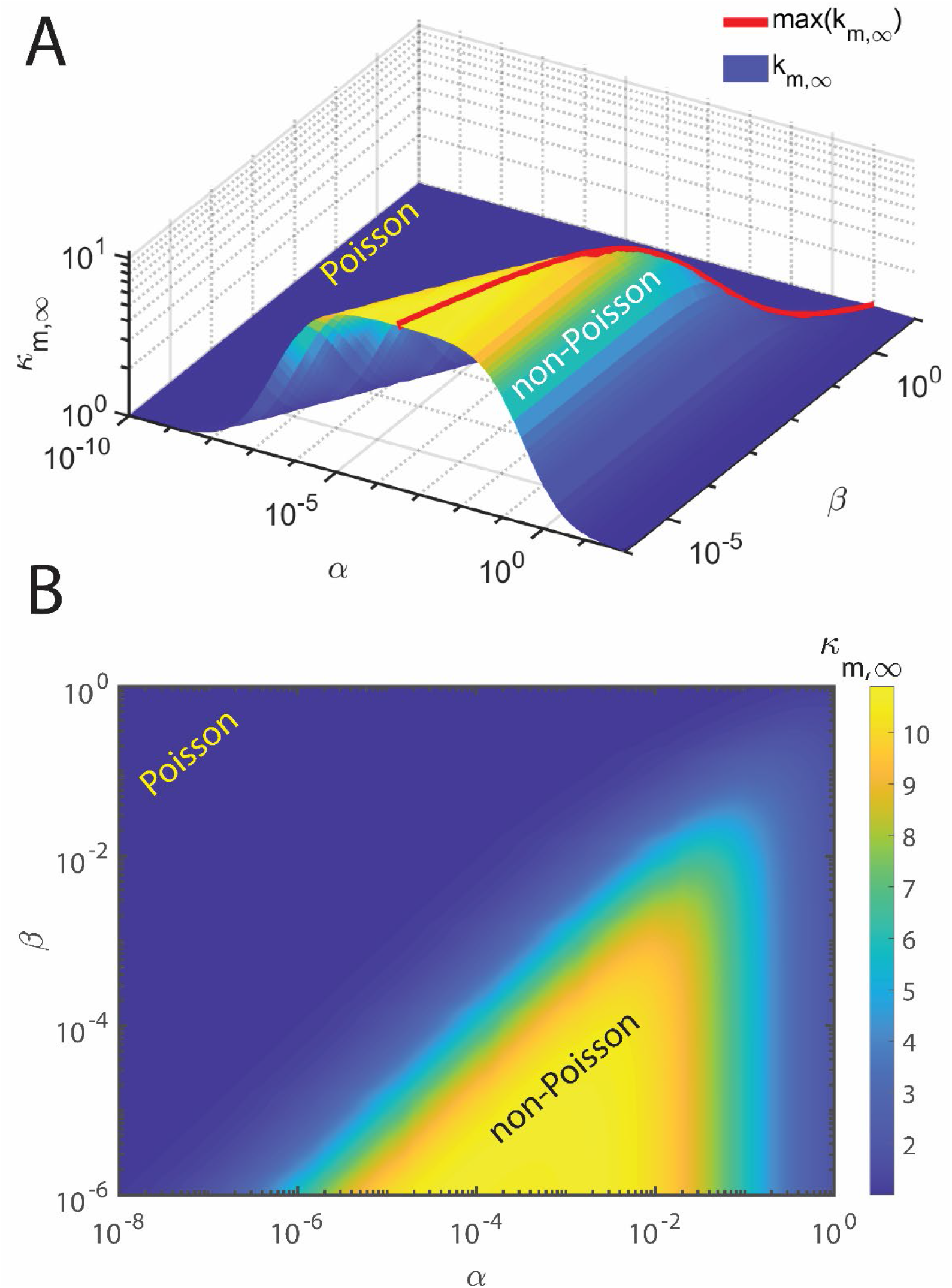
Variation of the steady state Fano factor *κ*_*m*,∞_ associated with the mRNA number fluctuations with respect to changes in the flipping rates *α* and *β* in the limit 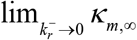 as given in **Eqs. 63**. Here the settings in panels **A** and **B** are 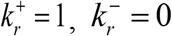 and *γ*_*r*_ = 0.1. Panel **B** is the contour plot of panel **A**. The optimum values of *α* at which *κ*_*m*,∞_ attains a maximum can be expressed as 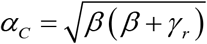. This can be obtained by solving [*∂κ*_*m*,∞_/*∂α*] = 0 for *α*. Red solid line in panel **A** is the maximum value of the steady state Fano factor which can be obtained by substituting *α* = *α_C_* in the expression of *κ*_*m*,∞_ as in **Eqs. 63**.

Generation of an individual mRNA transcript can be characterized by sequential microscopic transcription elongation rates 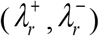 as shown in **Figs. 2**. We define the time that is required to generate a complete mRNA transcript of size *L* bp starting from zero number of bases as the first passage time (FPT). Inverse of this FPT is the transcription rate *ξ* associated with the respective mRNA transcript. Since FPT is a random variable across the population of transcription events, the resultant transcription rate will be a time dependent random variable in view of the mesoscopic transcription dynamics [44]. The average of FPTs is defined as the mean first passage time (MFPT). When there is a flipping across the on-off states, then the overall average transcription rate *k_r_* will be an inverse of the mean first passage time (*T_0_*, MFPT) as *k_r_* = [1 / *T*_0_]. When there is no flipping across the on-off states then one finds that 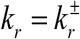 where 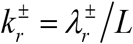 depending upon the transcription channel used to generate the mRNA transcript.

**Figs. 8** demonstrate how the statistical properties of FPTs such as mean, variance, Fano factor and coefficient of variation associated with the generation of a complete transcript varies with the on-off flipping rates. When we set *α* = *β* = *χ* and 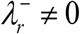, then there exists an optimum flipping rate *χ_C,T_* at which the variance and the Fano factor associated with the distribution of FPTs attain the maxima. This result is line with the prediction of **Eqs. A7** of **Appendix A**. Further, the Fano factor associated with the FPTs seems to be less than one throughout the iteration range of *χ*. This indicates the sub-Poisson type density function of FPTs. These results are demonstrated in **Figs. 8A-D**. When *α* ≠ *β* and 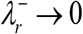, then the distribution of FPTs varies depending on the ratio (*α* / *β*) of the flipping rates.

**FIGURE 8.**
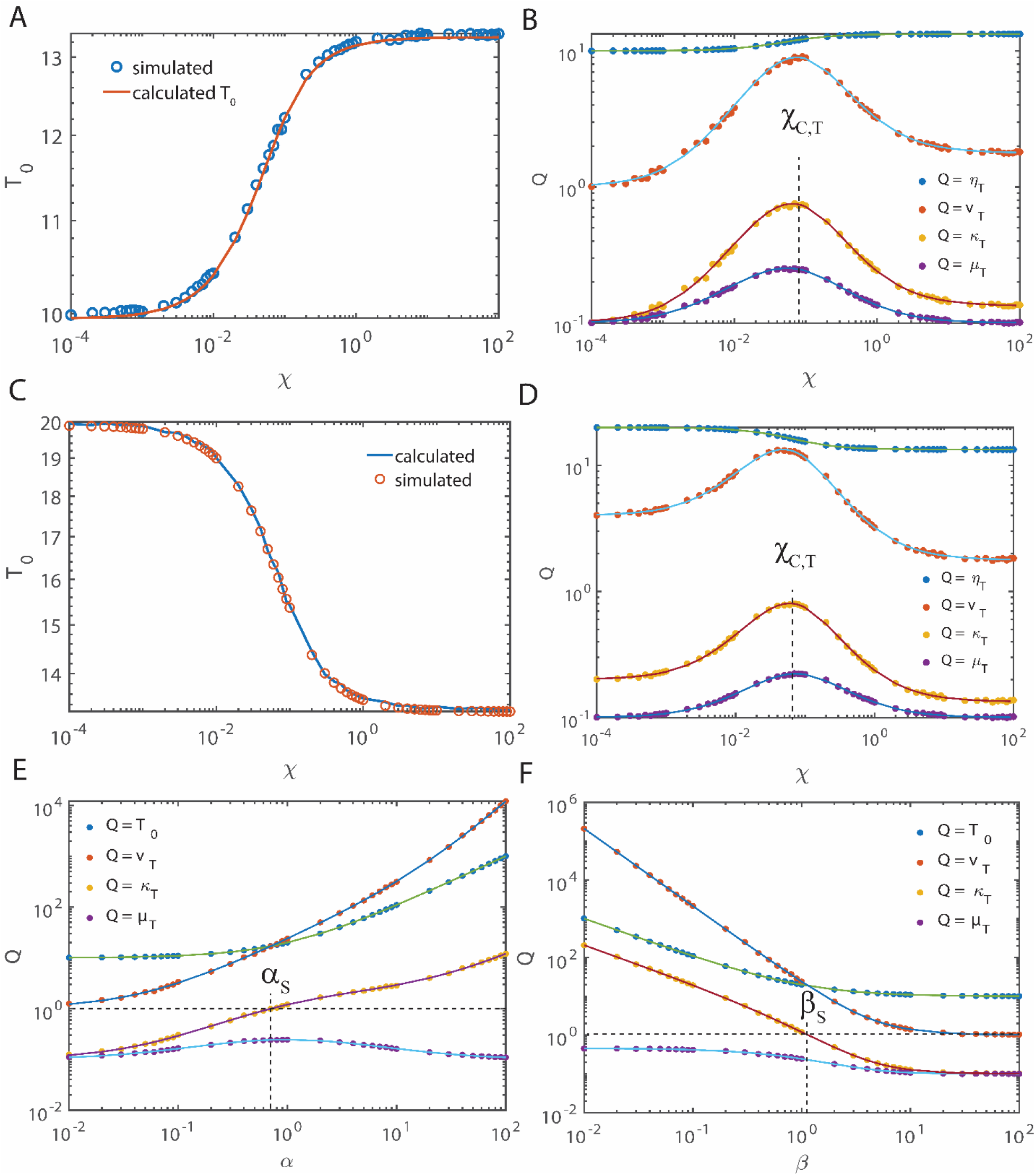
Dependence of the overall average transcription time on the on-off flipping rates. Here *T*_*0*_ is the mean first passage time associated with the formation of a complete mRNA transcript with size *n* = *L* bp starting from *n* = 0 bp where *n* is denotes dynamic number of transcribed bases. Filled or hollow circles are the simulated results and solid lines are the theoretical predictions computed using **Eqs. A3** (panels **E**, **F**) and **A7** (panels **A**-**D**) of **Appendix A**. By definition *k*_*r*_ = [1/*T*_*0*_] is the transcription rate. *T*_*0*_ was calculated using **Eqs. 39**. Hollow blue and red circles are the stochastic simulation results. In panels **A**-**D** the settings are 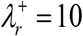, 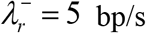, *α* = *β* = *χ* and *L* = 100 bp with 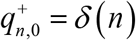 and 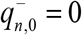 in **A** and **B**. In **C** and **D** 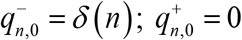. Here 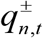 is the probability of finding *n* number transcribed bases in the respective on (+) and off (−) state channels. When the flipping rate becomes sufficiently large, then one finds the limiting value as 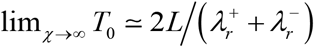. Panels **B** and **D** demonstrate the variation of the mean (*η*_*T*_), variance (*v*_*T*_), coefficient of variation (*μ*_*T*_) and Fano factor (*κ*_*T*_) of the first passage times associated with the generation of a complete mRNA transcript with respect to the flipping rate parameter *χ*. Clearly, there exists an optimum flipping rate *χ*_*C,T*_ at which the variance, Fano factor and coefficient of variation of the first passage times attain a maximum. Panels **E** and **F** demonstrate behavior of the overall MFPT when *α* ≠ *β* and 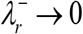. In this situation, we find 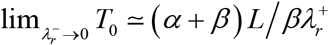. Settings in **E** and **F** are 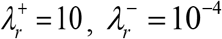 and *L* = 100. In panel **E**, *β* = 1 and in panel **F**, *α* = 1 with initial conditions 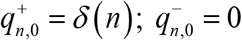. Statistical properties of FPTs were computed over 10^6^ number of trajectories. With these settings one finds that *β_S_* ~ 1.15 and *α_S_* ~ 0.7 in line with the prediction by **Eqs. 42**.

When (*α* / *β*) = 1, then the distribution of FPTs approximately follow a Poisson type density function with a Fano factor = 1. When (*α* / *β*) > 1, then the distribution of FPTs follow a super Poisson (over dispersed with a Fano factor > 1). When (*α* / *β*) < 1, then the distribution of FPTs follow a sub Poisson (under dispersed with a Fano factor < 1). These results are demonstrated in **Figs. 8E** and **F**. It is also interesting to note the limit 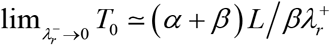. These results are consistent with the theoretical prediction by **Eqs. A3** of **Appendix A**. as we have shown in **Appendix A**. Various other limiting values of the MFPT (corresponding to the case 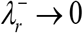) associated with the generation of a complete mRNA transcript are shown in **Figs. 9** (for the situation where *α* = *β* = *χ*) and **Figs. 10** (for the situation where *α* ≠ *β*). **Figs. 10** provides the computational proof for the various limiting values of the MFPT required to generate a full mRNA transcript as described in the **Appendix A**. Variation of the critical flipping rates (*α*_*S*_, *β*_*S*_) associated with distribution of FPTs with respect to (*α*, *β*) as described in **Eqs. 42** is demonstrated in **Figs. 10E** and **F**. Emergence of steady state bimodal type distribution of mRNA populations is demonstrated in **Figs. 11** along with a fitting to the experimental data digitized from Ref. [14].

**FIGURE 9.**
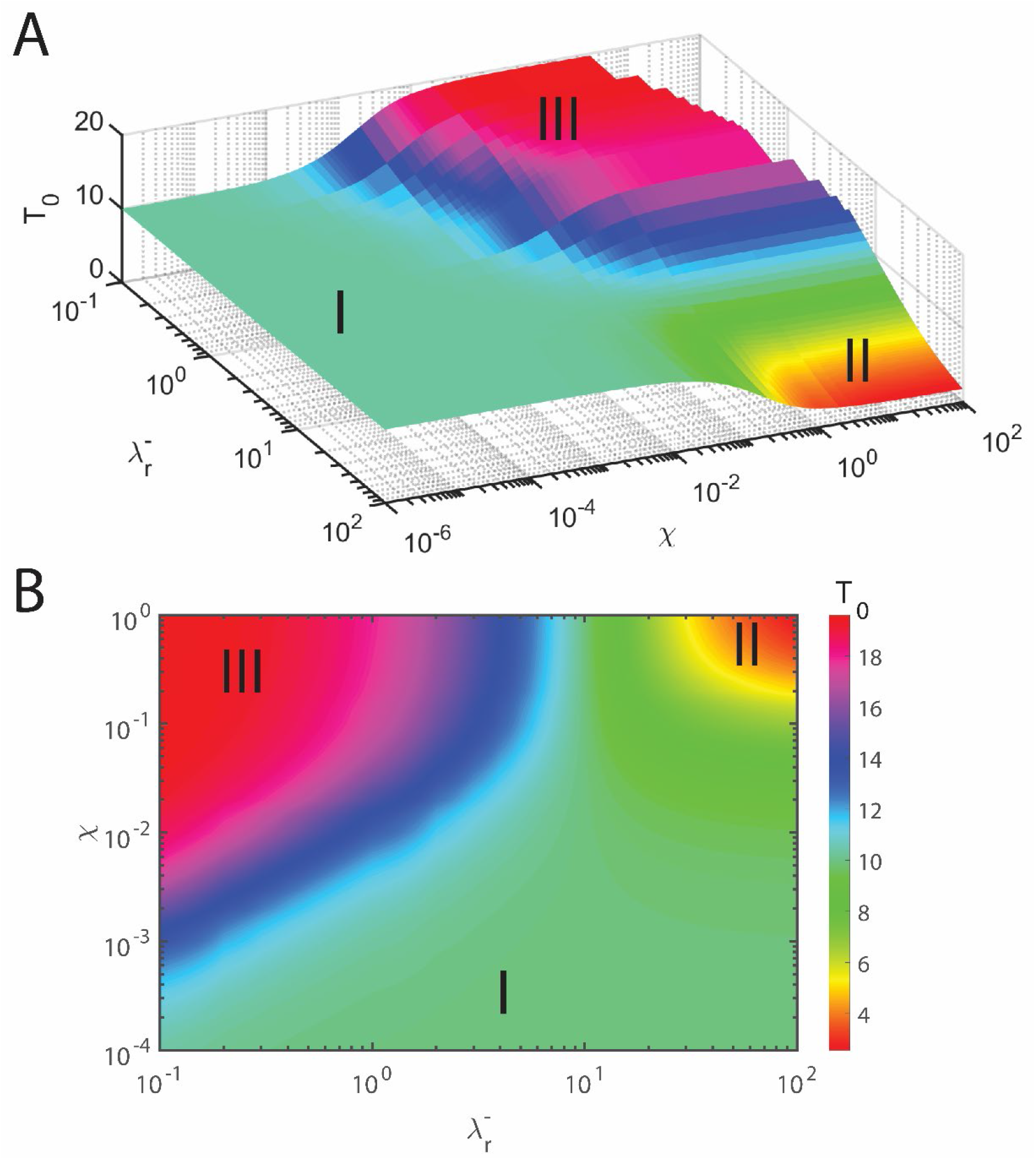
Various limiting values of the mean first passage time (MFPT, *T*_*0*_) associated with the generation of a complete mRNA transcript of size *L* bp. MFPT was computed using **Eqs. 44**. Here the settings in panel **A** are 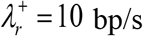 and *L* = 100 bp. When 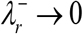 and *χ* → 0, then one finds that 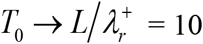 (**I**). When 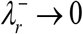 with finite *χ*, then one finds that 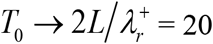 s (**III**). Hypothetically, when 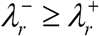 and *χ* → *∞*, then one finds that 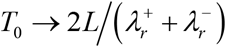 (**II**). Panel **B** is the contour plot of panel **A**.

**FIGURE 10.**
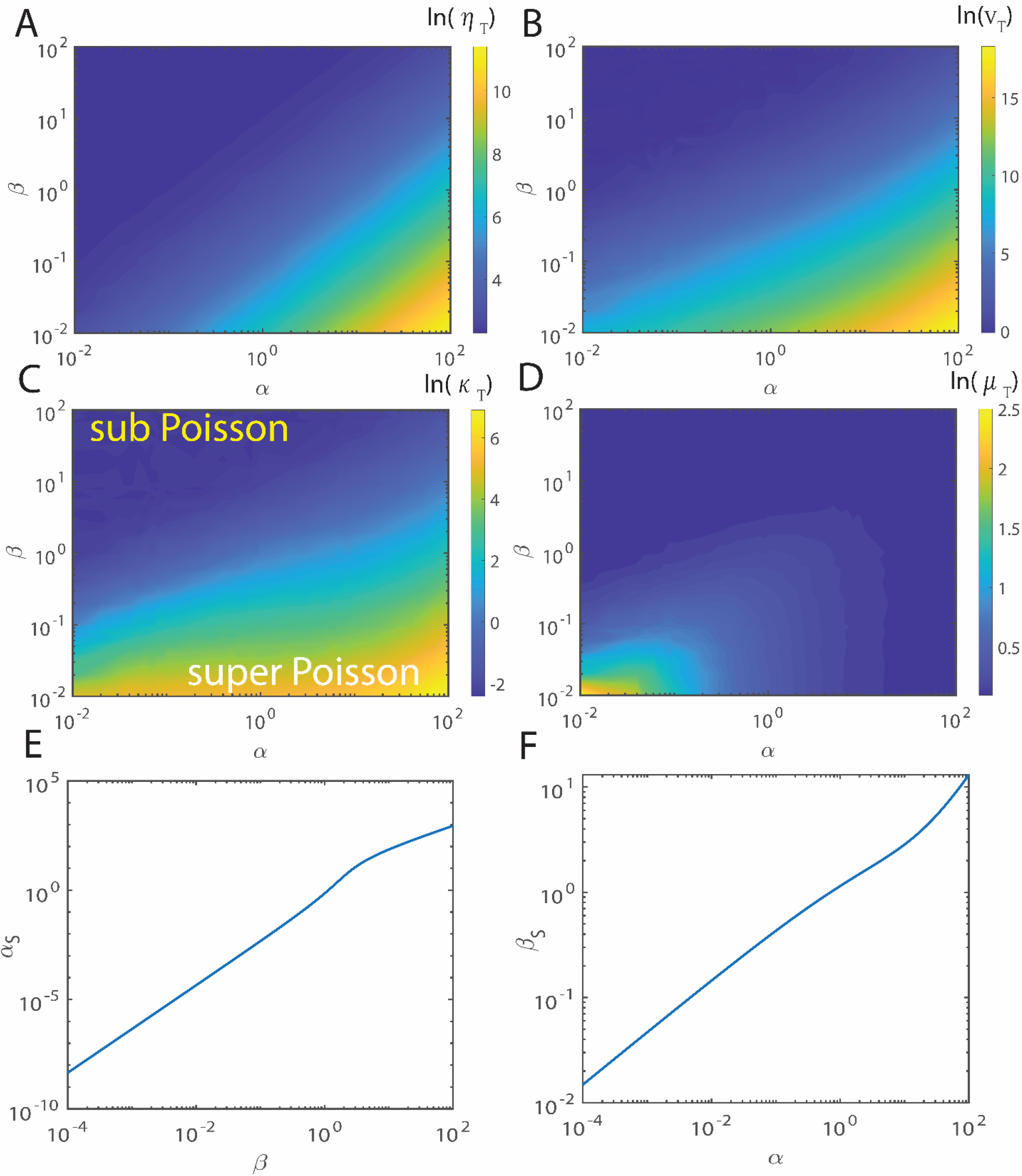
Statistical properties of the first passage times (FPTs) associated with the generation of a complete mRNA transcript of size *n* = *L* bp starting from *n* = 0. Settings are, 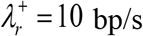 is the elongation rate in the on channel and *L* = 100 bp and 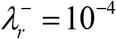 is the elongation rate of the off channel. As shown in **Appendix A**, one finds the limiting value of the mean of FPTs (MFPT, *T*_*0*_) as 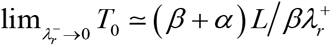. When *α* = 10^2^ and *β* = 10^−2^, then one finds the limiting value as 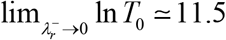. For *α* = 10^−2^ and *β* = 10^2^, one finds that 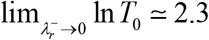 and so on. **A**. mean of FPTs (*η*_*T*_). **B**. Variance of FPTs (*v*_*T*_). **C**. Fano factor of FPTs (*κ*_*T*_). **D**. coefficient of variation of FPTs (*μ*_*T*_). Statistical properties of FPTs were computed over 10^6^ number of trajectories. In **E** and **F** (*α*_*S*_, *β*_*S*_) are the critical values of the flipping rates such that when *α* < *α*_*S*_ or *β* < *β*_*S*_, then the distribution of FPTs becomes sub-Poisson type. When *α* > *α*_*S*_ or *β* > *β*_*S*_, then the distribution of FPTs becomes super-Poisson as described in **Eqs. 42**. Here 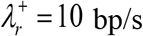. **E**. Variation of *α*_*S*_ with respect to *β* in the limit 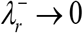. **F**. Variation of *β*_*S*_ with respect to *α* in the limit 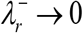.

**FIGURE 11.**
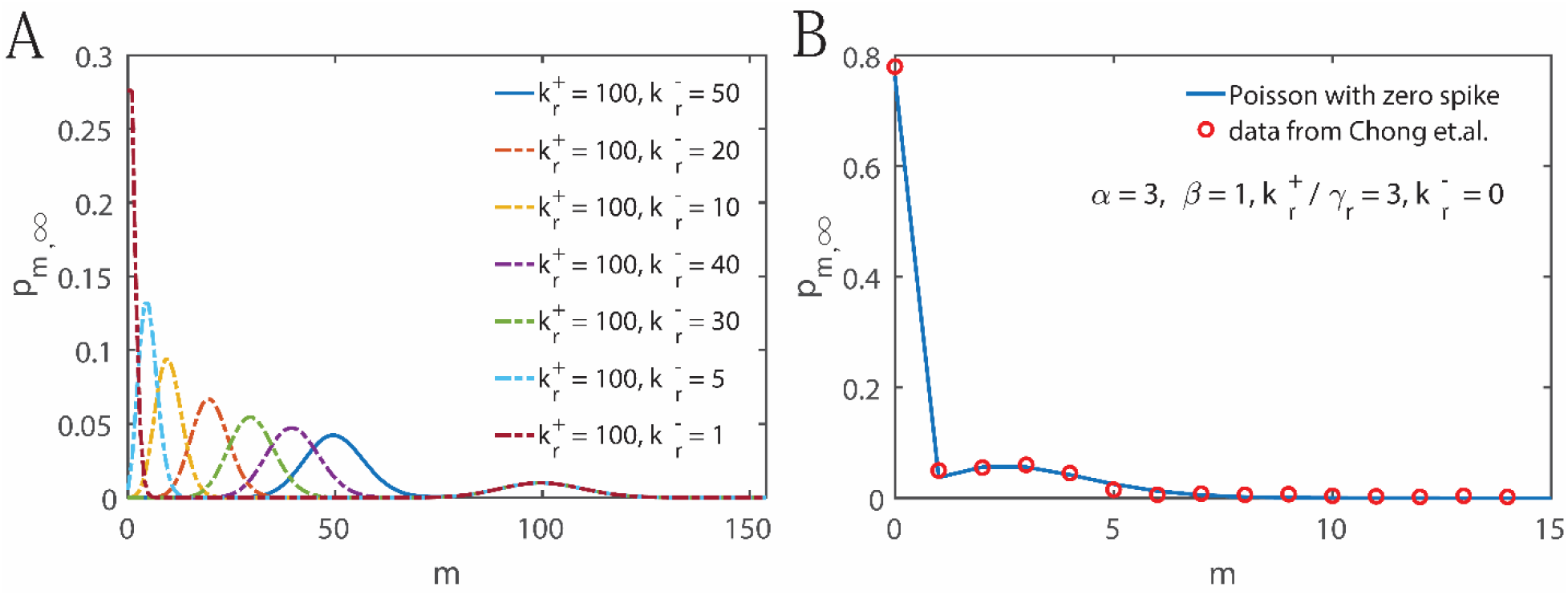
Emergence of the bimodal distribution functions associated with the mRNA number fluctuations. In panel **A** the settings are *α* = 3, *β* = 1 and *γ*_*r*_ = 1. In panel **B** the settings are *α* = 3, and *β* = 1. This represents the Poisson density function with zero spike as derived in **Eq. 78**. Hollow red circles are the digitized data points from Ref. [14].

One can broadly classify the models on transcription bursting into two state and multi state models [45]. In two state models, the promoter is assumed to flip across on and off states. Two state models generally produce a bimodal type density of the mRNA numbers. In the multistate models [19], there are several states across which the transcription process fluctuates. Multistate models generally result in the multimodal type density functions associated with the mRNA number fluctuations [46]. In all these models, the transcription rate *k_r_* will be assumed to be a constant. This assumption will work only when the timescale associated with the flipping across various states is much higher than the timescale associated with the generation of a complete mRNA transcript. It is generally assumed that the transcription machinery generates several mRNA transcripts (following the Poisson density function) in the on-state before flipping to the off-state e.g. the interrupted Poisson model developed in Ref. [47]. However close look at the underlying transcription mechanism reveals that the RNAP or RNA pol II performs several rounds of stall-continue type dynamics before generating a complete transcript. Therefore, the time that is required to generate a given transcript will be a random variable. This means that the resultant transcription rate *ξ* (which is the inverse of the first passage time required to generate a complete mRNA transcript) will be a random variable.

The overall average transcription rate will be defined as *k_r_* = ∫ *ξp* (*ξ*) *dξ* = ⟨*ξ*⟩ where *p* (*ξ*) is the probability density function associated with the distribution of transcription rates. Depending on the timescale of on-off state flipping one can consider two different scenarios viz. (**a**) when the timescale of on-off flipping is much longer than the timescale associated with the generation of a complete mRNA transcript, then the transcription rate will be homogenous within a given on-state period but varies from burst to burst. This can be approximately described as a static disorder in the transcription rates. (**b**) When the timescale of flipping is much shorter than the timescale associated with the mRNA synthesis, then the transcription rate varies from mRNA to mRNA. This can be described as a dynamical disorder in the transcription rates [44, 48, 49]. Our mean first passage time calculations suggested that when the rate of off-state transcription channel is zero i.e. 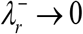, then the overall average transcription rate will be transformed as 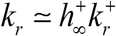 where 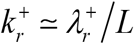 and 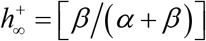 is the stationary state probability of finding the transcription machinery in the on-state (**Appendix A** and **Eqs. B8** of **Appendix B** or sufficiently large mRNA length *L*) in the presence of flipping across the on-off states. This means that the transcription “rate constant” will be a function the on-off flipping rates. Comparison of this quantity with the steady state and time dependent solutions of **Eqs. 1** clearly suggested that **Eqs. 1** is accurate enough (this can be inferred from the expression of *η*_*m*,*t*_ in **Eqs. 11** and *η*_*m*,∞_ of **Eqs. 58**) to capture the inhomogeneity of the resultant transcription rate *ξ* at sufficiently large mRNA lengths. However, **Eqs. 1** cannot explain the origin of over-dispersion in the mRNA numbers. Our detailed study reveals that such over-dispersion mainly originates from the non-Poisson type distribution of FPTs associated with the elongation of individual mRNA transcripts especially when the on-off flipping rates are such that (*α* / *β*) > 1.

## CONCLUSION

Transcription bursting is essential to generate variation among the individuals of a given population. The mechanism of bursting comprises of at least three sub-processes with different timescale regimes viz. flipping dynamics across the on-off state transcription elongation channels, microscopic transcription elongation events and the mesoscopic transcription dynamics along with the mRNA recycling. Flipping dynamics across the on-off states is similar to the tossing of a coin. When the flipping dynamics is combined with the microscopic elongation events, then the distribution of resultant transcription rates will be over-dispersed. This in turn reflects as the over-dispersed non-Poisson type distribution of mRNA numbers.

Our detailed calculations show that there exist optimum flipping rates (*α_C_*, *β_C_*) at which the stationary state Fano factor and variance associated with the mRNA numbers attain maxima. These optimum points are connected via 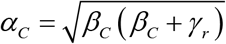. Here *α* is the rate of flipping from the on-state to the off-state and *β* is the rate of flipping from the off-state to the on-state of transcription and *γr* is the decay rate of mRNA. When *α* = *β* = *χ*, then there exist optimum flipping rates at which the non-stationary Fano factor and variance attain maxima. Here 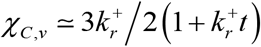 (where 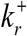 is the rate of transcription through the on-state channel) is the optimum flipping rate at which the variance of mRNA numbers attains a maximum and *χ*_*C,κ*_ ≃ 1.72 / *t* is the optimum flipping rate at which the Fano factor attains a maximum. These optimum points reduce to zero when *t* → ∞.

Close look at the transcription mechanism reveals that the RNA polymerase enzyme complex performs several rounds of stall-continue type dynamics before generating a complete mRNA transcript. Based on this observation we model the transcription event as the stochastic trajectory taken by the transcription machinery across these on-off state elongation channels. Each transcript follows different trajectory. The total time taken by a given trajectory is the first passage time (FPT). Inverse of this FPT is the resultant transcription rate *ξ* associated with the particular mRNA. Therefore, the time that is required to generate a given mRNA transcript will be a random variable. This means that the resultant transcription rate *ξ* will be a random variable. The overall average transcription rate will be the ensemble average *k_r_* = ⟨*ξ*⟩ which is equal to the inverse of the mean first passage time required to generate a complete mRNA. For a stall-continue type dynamics of RNA polymerase, the overall average transcription rate can be expressed as 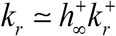 where 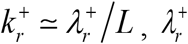 is the microscopic transcription elongation rate on the on-state channel and *L* is the length of complete mRNA and 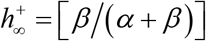 is the stationary state probability of finding the transcription machinery in the on-state.

**Table 1.**
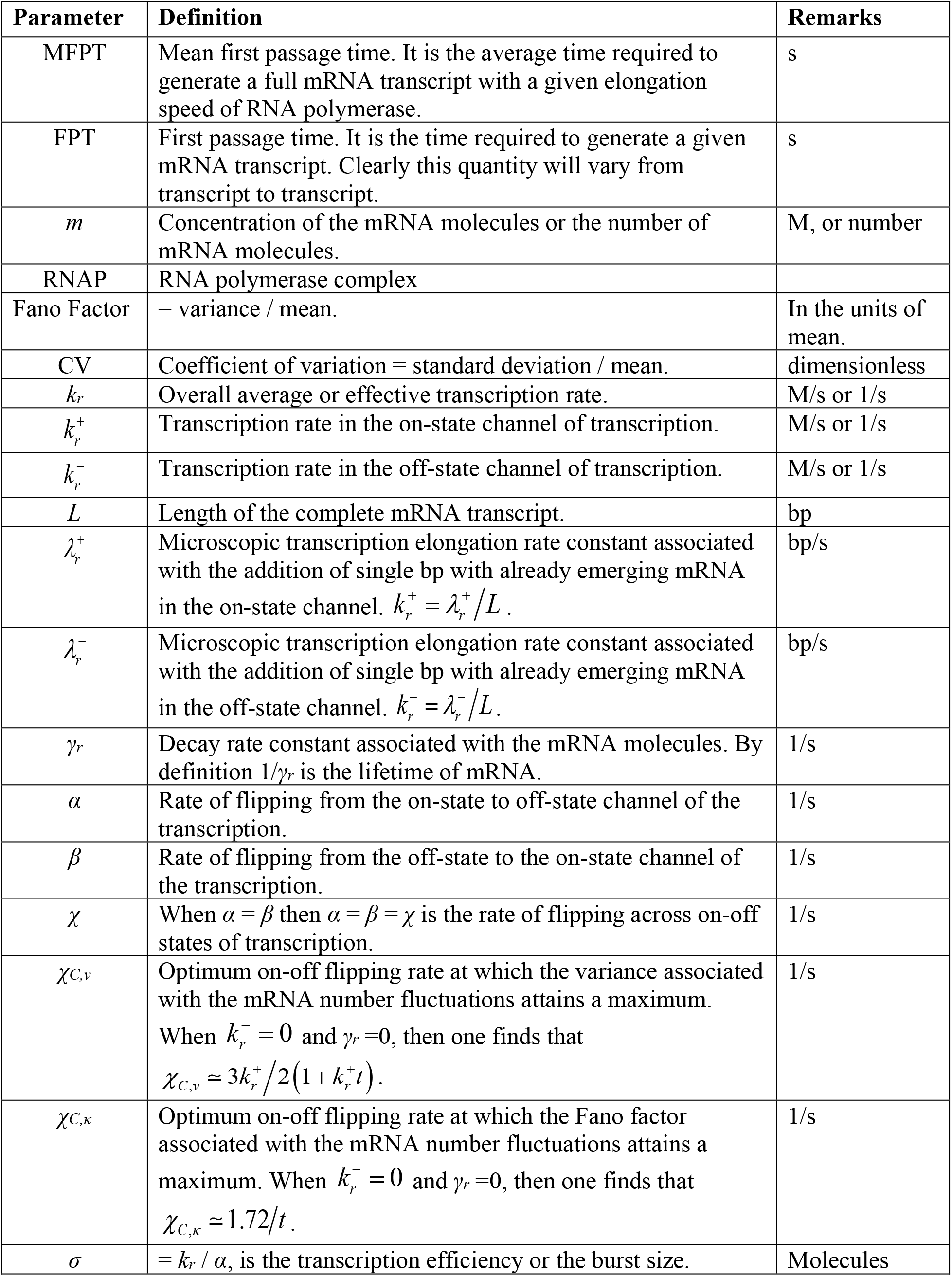
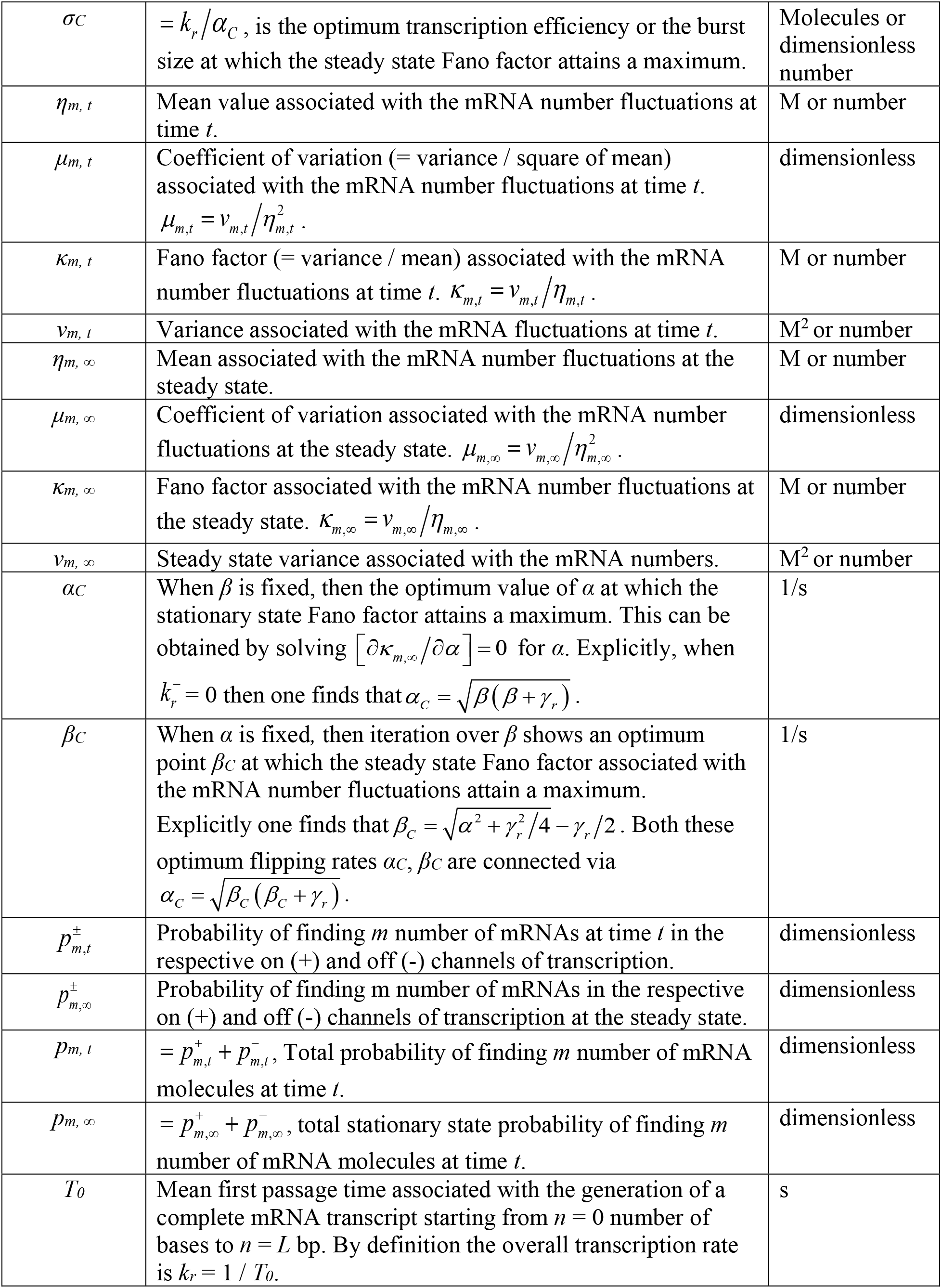
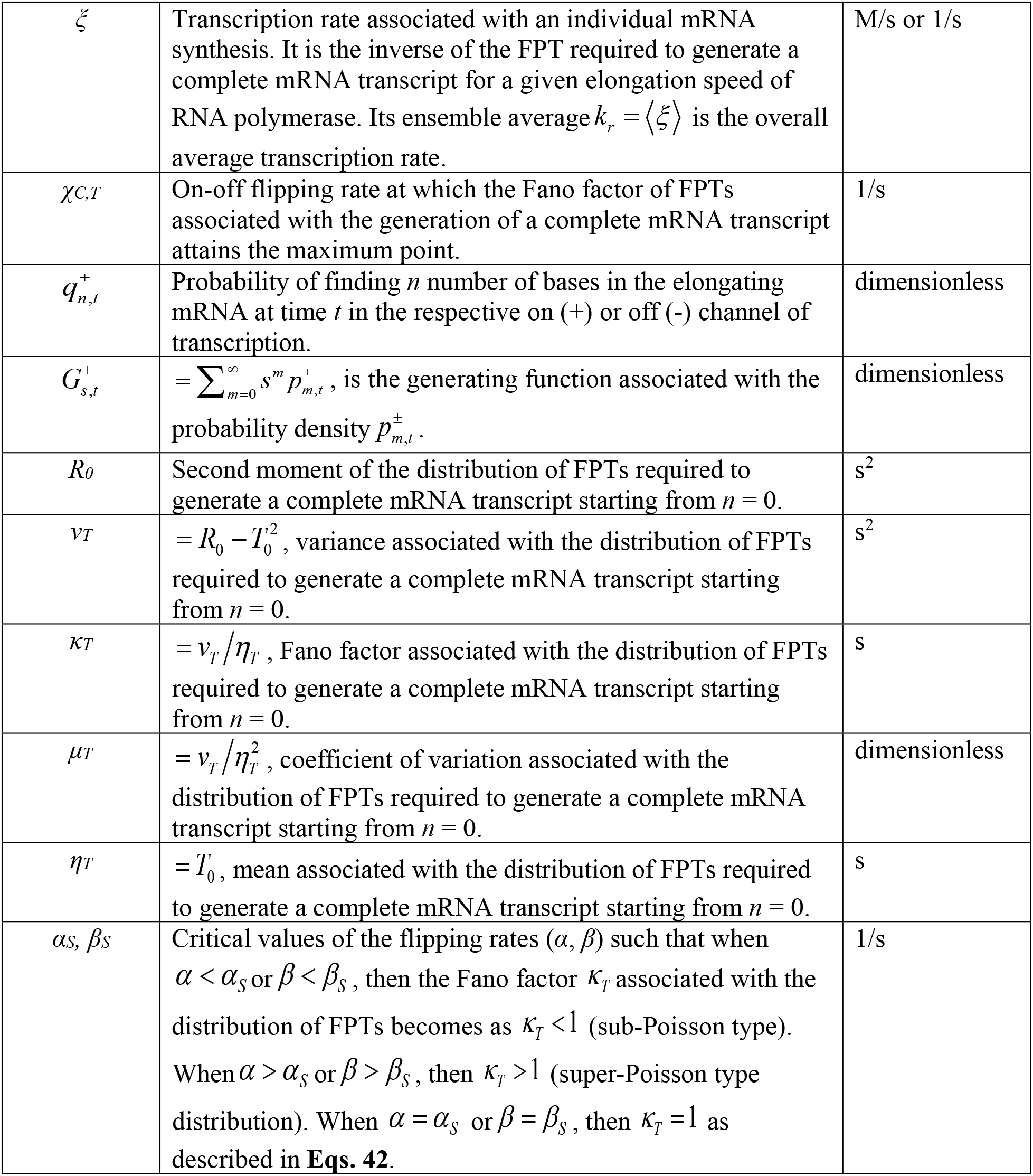
List of symbols and variables used in the main text.

## APPENDIX A

Consider the following backward type coupled master equations (**Eqs. 40** of the main text) corresponding to the second moments of the first passage times (FPTs) required to generate a complete mRNA transcript starting from an arbitrary *n* size of mRNA [37, 39].

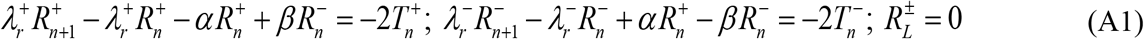

In this equation, the first moments of the FPTs are defined as follows (as described in **Eqs. 42** of the main text).

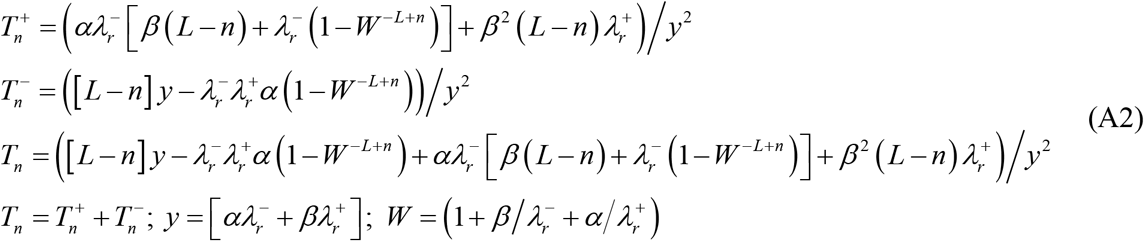

The set of difference **Eqs. A1** is exactly solvable using standard summing technique [37] and the final expression corresponding to the variance (*v_T_*), Fano factor (*κ*_*T*_) and coefficient of variation (*μ*_*T*_) of FPTs associated with the generation of a complete mRNA transcript of length *L* starting from *n* = 0 can be written as follows.

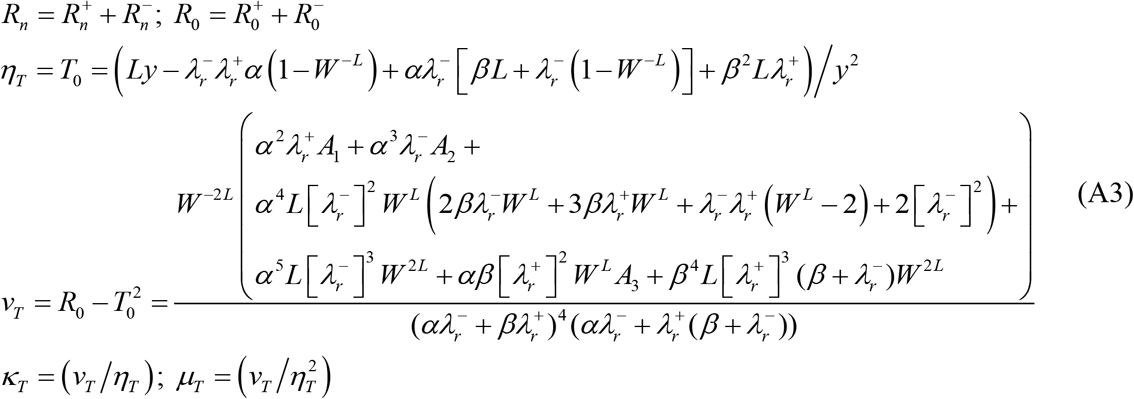

Here *y* and *W* are defined as in **Eqs. A2** and various other terms *A_1_*, *A_2_* and *A_3_* in **Eqs. A3** are defined as follows.

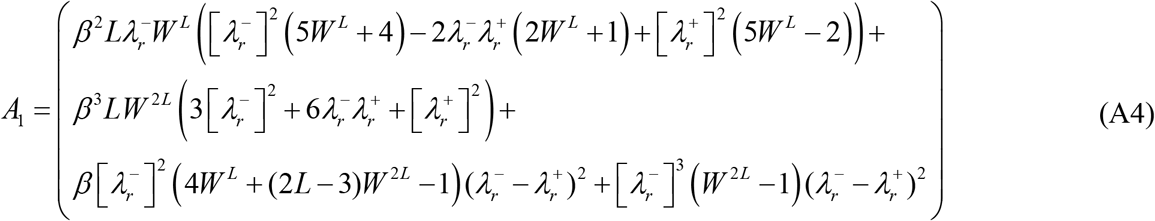

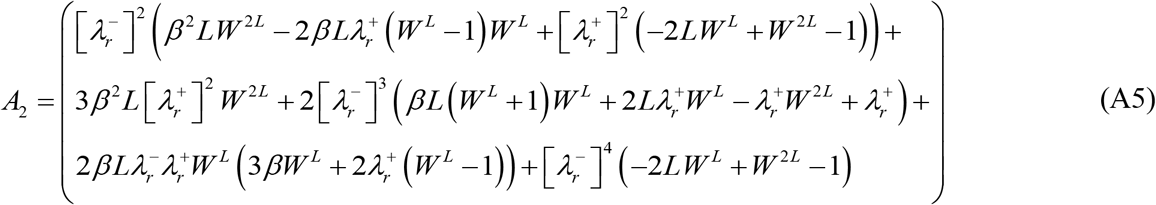

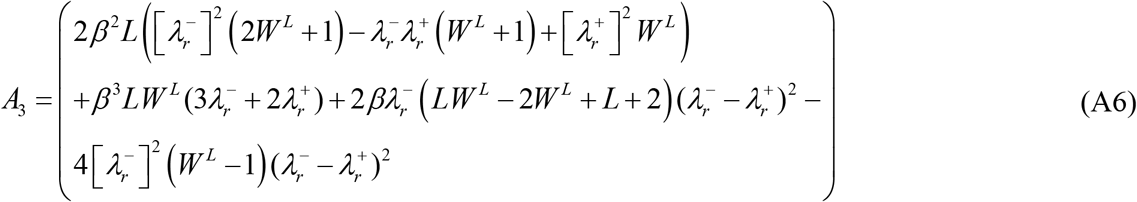

In **Eqs. A4**-**A6**, *W* is defined as in **Eqs. A2**. When *α* = *β* = *χ*, then the expression corresponding to the variance of FPTs given in **Eqs. A3** simplifies to the following form.

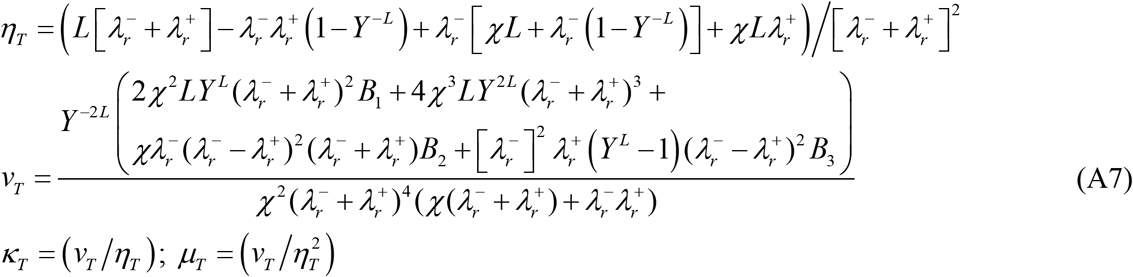

The terms *Y*, *B_1_*, *B_2_* and *B_3_* in **Eqs. A7** are defined as follows.

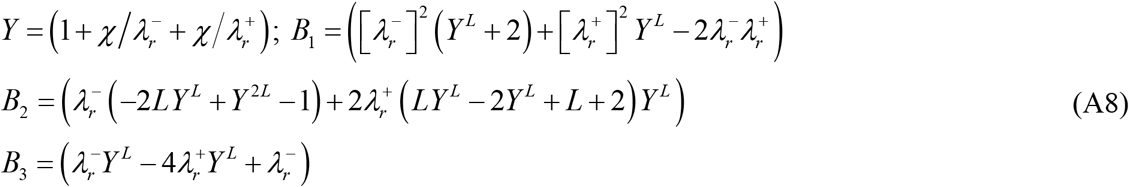

Numerical analysis of **Eqs. A7** reveals the existence of maximum *v_T_*, *κ*_*T*_ and *μ*_*T*_ at the optimum flipping rate *χ_C,T_* which is demonstrated in **Figs. 8B** and **D**. **Eqs. A2** clearly suggest that when the microscopic transcription elongation rate associated with the off-state channel is close to zero i.e. 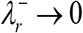 then the overall average transcription rate scales with the on-off flipping rates as 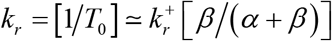 where 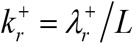 is the overall transcription rate of the pure on-state channel and 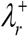 is the elongation rate of the on-state channel.

## APPENDIX B

Let us consider the uncoupled on-off state transcription elongation channels. When there is no flipping across the on and off state channels, then **Eqs. 1** will be uncoupled as follows.

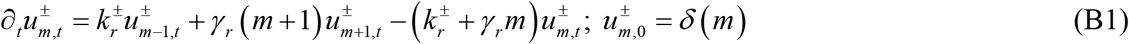

Here 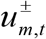 is the probability of finding *m* number of mRNAs at time *t* in the respective independent transcription channels. Upon defining the generating functions as 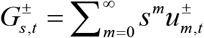 one can transform **Eqs. B1** into the following partial differential equations.

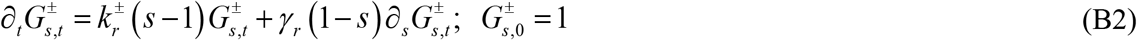

Upon solving **Eqs. B2** for the for the appropriate boundary conditions one finds the expression for the generating functions as follows.

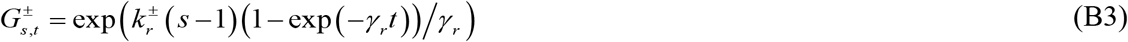

Using this generating function one can derive various statistical properties of the mRNA number fluctuations corresponding to on-off channels as follows.

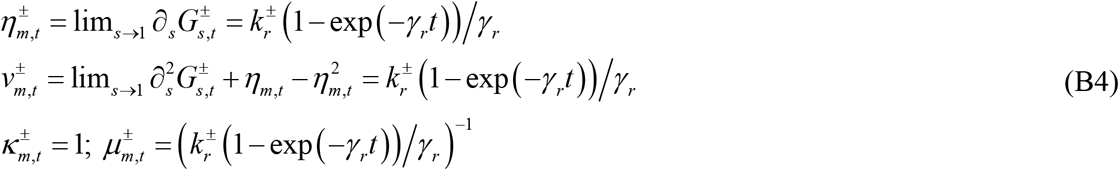

Upon expanding the generating function in to a Macularin series with respect to variable *s* and then setting *s* = 1, one finally obtains the probability density functions as follows.

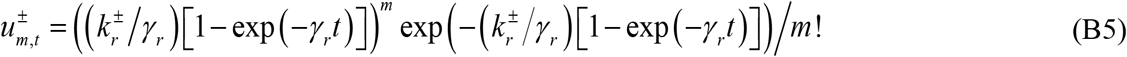

Using **Eqs. B5** one can directly obtain the steady state probability density functions as follows.

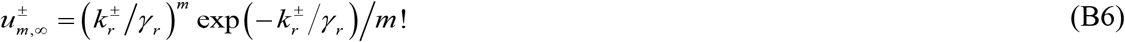

When the on-off state flipping is uncoupled from the transcription elongation, then the flipping dynamics can be described by the following set of coupled differential equations.

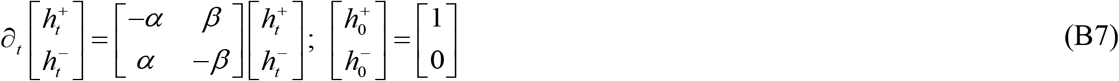

Here we have the normalization condition 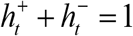. Upon solving **Eqs. B7** with the appropriate initial conditions, one obtains the expression for the time dependent probability 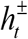 of finding the transcription state in the respective on-off channels as follows.

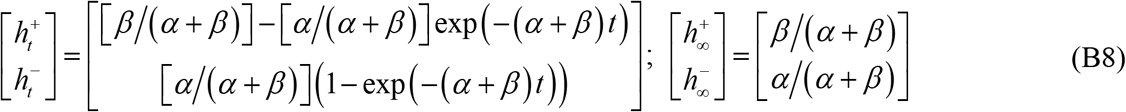

Upon combining **Eqs. B5** with **Eqs. B8**, one can derive 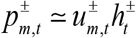, which is the probability of finding *m* number of mRNAs in the respective on-off state channels. Explicitly one can write the following expression.

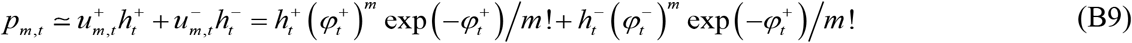

In this equation, we have defined the function 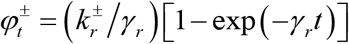. When the timescale associated with the on-off state flipping dynamics is much lower than the timescale associated with the mRNA synthesis and decay dynamics so that (*α* + *β*) ≫ *γ*_*r*_, then **Eq. B9** reduces to 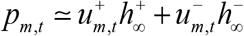. Under complete steady state conditions 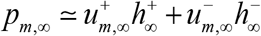 and one obtains the following bimodal Poisson type expression.

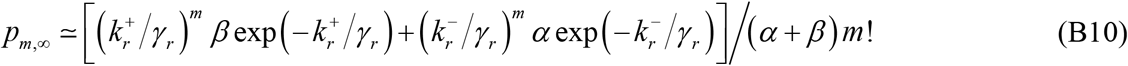

When the transcription rate associated with the off-state channel tends toward zero, then one recovers the Poisson density function with zero spike [14] as follows.

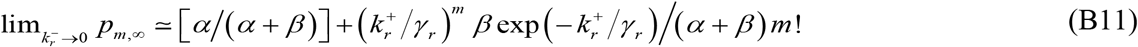

## Notes

#### Summary of Updates

solution to higher order moments of FPTs were included along with simulation results.

